# Disrupting Notch signalling by a small molecule inhibiting dihydroorotate dehydrogenase activity

**DOI:** 10.1101/2025.09.29.679158

**Authors:** Eike-Benjamin Braune, Dirk Wienke, Anita Seshire, Timo Heinrich, Martin Haraldsson, Sonia Lain, Urban Lendahl

## Abstract

The Notch signalling pathway is highly evolutionarily conserved and regulates differentiation and homeostasis in most organs. Given the critical role of Notch signalling for normal development, dysregulated Notch signalling is frequently linked to pathogenesis of disease and cancer. Hence, developing Notch-targeting therapeutics is warranted but has been challenging and Notch inhibitors have not yet reached broad clinical use. In this report, we identify potential Notch inhibitors, using a novel cell-based Notch reporter system for unbiased screening of compounds reducing Notch signalling. A library of 37.966 small organic compounds was screened for inhibitor candidates, followed by a counter screen to eliminate γ-secretase inhibitor-like compounds and an orthogonal screen based on the role of Notch signalling in myogenic differentiation. This triage led to the identification of five Notch inhibitor candidate hits with different chemical backbones and unrelated to previous Notch antagonists. One candidate hit showed structural similarities to dihydroorotate dehydrogenase (DHODH) inhibitors, and we provide evidence that inhibition of DHODH activity reduces Notch signalling. In conclusion, our data support the notion that DHODH inhibition may be an interesting avenue to explore for the development of novel Notch inhibitors.

## Introduction

The Notch signalling pathway is a cell-cell communication system required for differentiation and homeostasis of most tissues and organs. Notch receptors and ligands are highly evolutionarily conserved, and Notch signalling operates in most, if not all, multicellular organisms (Bray, 2016; Siebel & Lendahl, 2017). In most tissues, Notch signalling promotes an undifferentiated cell state and blocks differentiation, while in some organs, such as the skin, Notch signalling drives differentiation.

Mechanistically, Notch signalling is initiated when a transmembrane Notch receptor interacts with a transmembrane Notch ligand of the Jagged or Delta-like type presented on a juxtaposed cell (Figure 1A). Ligand-receptor interaction leads to proteolytic processing of the Notch receptor by ADAM10 at the extracellular side close to the plasma membrane (referred to as S2-cleavage). S2-cleavage is rapidly followed by S3-cleavage, which is executed by the γ-secretase complex at the intracellular surface of the plasma membrane or in endosomes (Figure 1A). S3-cleavage liberates the C-terminal portion of the Notch receptor, i.e., the Notch intracellular domain (Notch ICD), which translocates to the cell nucleus. In the cell nucleus, Notch ICD interacts with the DNA-binding protein CSL and an adaptor protein MAML. The ternary Notch ICD/MAML/CSL complex regulates transcription of downstream genes, including *Nrarp*, *Hes* and *Hey* (Figure 1A).

**Figure 1:**
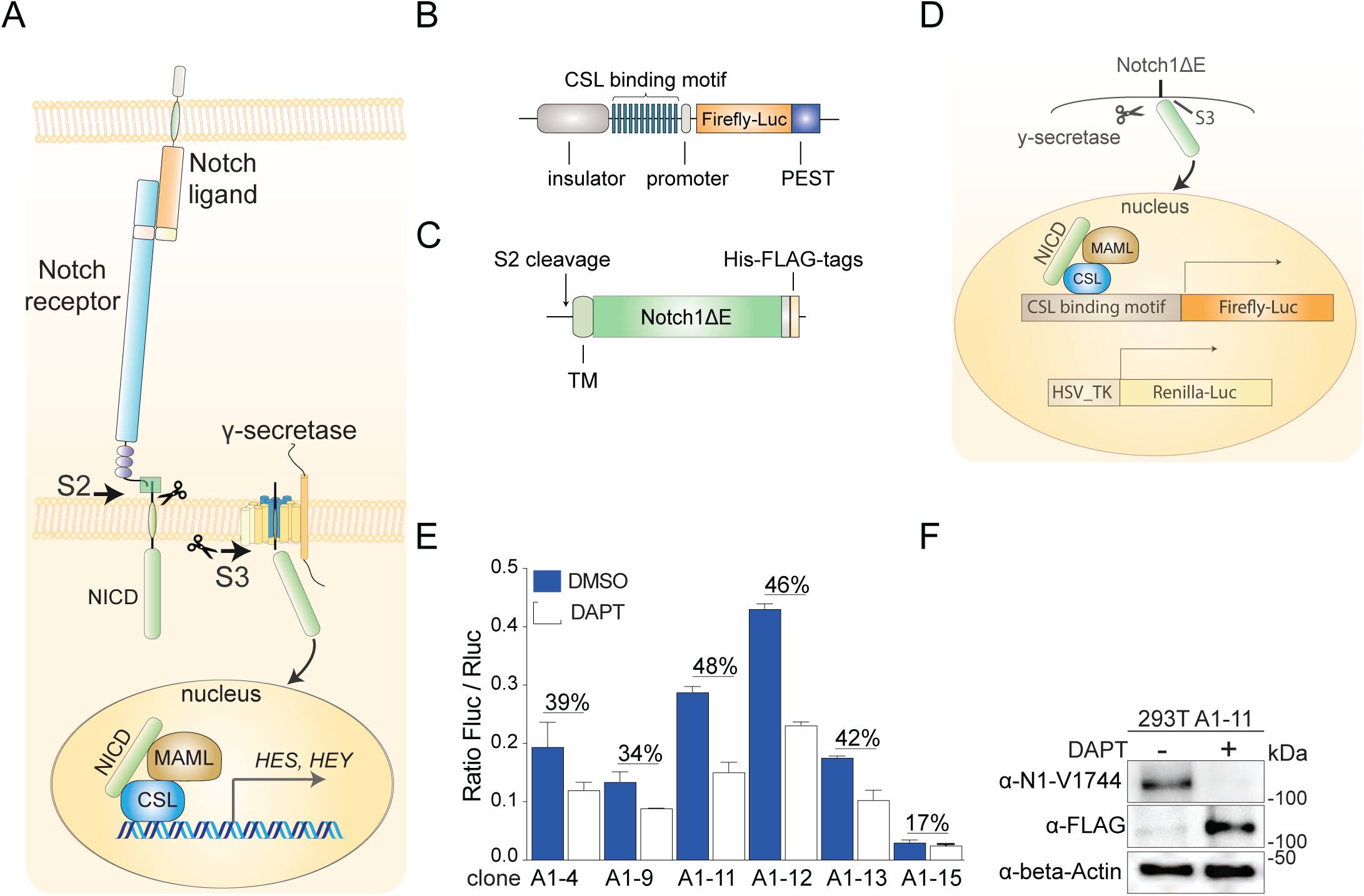
A novel Notch reporter system. **(A)** Schematic overview of the Notch signalling pathway. Arrows mark the S2 and S3 cleavage sites, and the ternary Notch ICD/MAML/CSL transcriptional complex in the nucleus is depicted. **(B)** Schematic depiction of the novel Notch reporter construct. **(C)** The Notch1DE construct used to generate enhanced ligand-independent Notch signalling in the reporter cell line. The S2 cleavage site, transmembrane domain (TM) and C-terminal protein tags are indicated. **(D)** Schematic depiction of the reporter cell line, containing the Notch1DE construct, the CSL-firefly luciferase reporter to read out Notch signalling and the constitutively activated HSV-TK renilla-luciferase as an internal control for luciferase activity. **(E)** Analysis of the levels of firefly/renilla luciferase activity in different clones of the HEK293T reporter cell line. **(F)** Western blot analysis of Notch1DE in clone #A1-11 in the presence or absence of DAPT, visualized by an anti-FLAG (a-FLAG) antibody or an antibody recognizing the S3-cleaved form of NOTCH1 (a-N1-V1744), as indicated. beta-Actin was used as loading control.

In accordance with its importance for normal cellular differentiation, aberrant Notch signalling is observed in disease and cancer. Several monogenic diseases are caused by mutations in genes in the Notch signalling pathway, including a number of heart diseases, the multiorgan disease Alagille syndrome, the stroke and dementia syndrome CADASIL and the connective tissue disorder Hajdu-Cheney disease (Siebel & Lendahl, 2017). A considerable number of cancers, including acute lymphoblastic T-cell leukemia (T-ALL), adenoid cystic carcinoma (ACC), triple negative breast cancer (TNBC) and non-small cell lung cancer (NSCLC) are linked to hyperactive Notch signalling. There are however also tumour forms in which Notch serves as a tumour suppressor, including small cell lung cancer (SCLC), bladder cancer and squamous cell carcinoma (SCC) (Aster et al., 2017; Braune et al., 2018; Braune & Lendahl, 2016).

The observation that elevated Notch signalling, caused by Notch receptor gain-of-function mutations or hyperactivation of Notch signalling as an indirect consequence of other mutations, is found in several cancers and diseases has motivated efforts to develop Notch-targeting therapies. There have been several attempts to generate Notch inhibitors, and there are > 400 clinical trials with preclinical Notch inhibitors reported at clincialtrial.com. GSIs, which block the activity of the γ-secretase complex and thus S3 cleavage of all Notch receptors, have however caused severe side effects, e.g., goblet cell metaplasia, immune suppression and skin cancer, and it is therefore of interest to explore alternative, non-GSI-based inhibitors for future clinical use. Other types of small molecule inhibitors have been developed, including RIN1 (Hurtado et al., 2019), CB-103 (Lehal et al., 2020), NADI-351 (Alvarez-Trotta et al., 2021) and Z271-0326 (Diluvio et al., 2023), which disrupt the Notch transcriptional complex, and CB-103 is currently investigated in phase 2 clinical trials. Compounds interfering with the Golgi apparatus or Notch posttranslational modification are also being explored (Krämer et al., 2013; Lu et al., 2018). Receptor-specific antibodies that block Notch receptor-ligand interaction or receptor processing represent an alternative approach which has shown promise, and antibodies that specifically block each of the four NOTCH receptors or Notch ligands are currently available (Chung et al., 2017; Lafkas et al., 2015; Tran et al., 2013; Wei et al., 2020; Y. Wu et al., 2010). These reagents have the advantage that they specifically target the activity of an individual receptor, but long-term use in preclinical animal models is associated with side effects (Yan et al., 2010). There are to date no Notch inhibitors in routine clinical use (Andersson & Lendahl, 2014; Majumder et al., 2021), except for the γ-secretase inhibitor (GSI) nirogacestat, which recently received FDA priority review for soft-tissue desmoid tumours with no FDA-approved treatments (Keam, 2024).

In the light of the outlined limitations of current Notch-targeted approaches, new approaches are warranted to identify alternative Notch-directed therapies. In this report, we have taken an unbiased approach to identify novel Notch inhibitors based on a novel reporter assay recording signalling immediately downstream of the Notch receptor. We report the identification of five Notch inhibitor candidates with distinct chemical backbones, one of which constitutes a dihydroorotate dehydrogenase (DHODH) inhibitor. This identifies a role for DHODH in regulation of Notch signalling and our data suggest that DHODH inhibition may be an interesting avenue in the search for new ways to inhibit Notch signalling.

## Results

### A novel Notch reporter assay recording immediate downstream Notch signalling

To identify novel Notch inhibitors in an unbiased manner, we generated a novel Notch reporter construct, which records the level of Notch signalling immediately downstream of receptor activation rather than further downstream in the signalling cascade, for example at the level of Hes and Hey proteins (Figure 1B). The reporter construct is based on a set of 12 CSL-binding sites linked to a firefly luciferase gene with a PEST domain attached C-terminally, to decrease reporter gene half-life and increase the temporal dynamics of the assay (Figure 1B). The reporter construct carries piggy-bac sequences at both ends of the promoter-reporter cassette as well as cSH4 insulator elements to reduce transcriptional noise (S. Wu et al., 2006) and was transfected into 293T cells. To achieve a robust level of Notch receptor activity in the 293T reporter cell line and to confine the search for inhibitors to compounds affecting the receptor function and eliminating the need for ligand activation, a truncated ligand-independent version of the Notch1 receptor, mimicking an S2-cleaved Notch1 receptor (Notch1ΔE) and spontaneously activated by S3-cleavage (Chapman et al., 2006), was introduced into the cell line carrying the reporter construct (Figure 1C). In addition, an HSV-TK-renilla luciferase plasmid was introduced into the cell line as an internal control for normalisation of the reporter expression (Figure 1D).

A set of clones containing the reporter construct, the truncated Notch1 receptor and the renilla luciferase control were analysed for response to the GSI tert-Butyl (S)-{(2S)-2-[2-(3,5-difluorophenyl)acetamido]propanamido}phenylacetate) (DAPT), and the clone with strongest DAPT-mediated signal reduction was selected (clone A1-11; hereafter referred to as clone A) (Figure 1E). Western blot analysis revealed that the majority of Notch1ΔE in clone A was spontaneously S3-processed, as expected, while processing was almost completely abolished when DAPT was supplemented to the cells (Figure 1F). We next established the cell-based reporter system in a 384-well format, and both the firefly and renilla luciferase reporter activities were stable over time (>1h), produced Z-scores >0.5 and the renilla signal was not affected by DAPT, as desired (Supplemental Figure 1A; for evaluation of DSMO and DAPT effects see Supplemental Figure 1B,C). Together, these data suggest that the novel reporter assay can dynamically read out Notch signalling and functions in a 384-well format to enable high throughput screening (HTS) of inhibitors.

### Screening of a small compound library for novel Notch inhibitors

To identify novel Notch inhibitors, we opted for screening a focused library consisting of 37.966 small molecule compounds, selected to represent a broad structural diversity, while keeping a certain depth to allow crude structure–activity relationship studies (Beresini et al., 2014). The selection covered a large chemical space and was biased towards lead-like and drug-like profiles regarding molecular weight, hydrogen bond donors/acceptors and lipophilicity of compounds. The library furthermore included a nucleoside set from Berry & Associates as well as 1.200 compounds from the Prestwick collection of small compounds, which contains several EMA- and FDA-approved compounds and has specially been designed to increase potential high-quality hits (https://www.prestwickchemical.com/). As controls, we included a set of Notch inhibitors and GSIs. The small molecule Notch inhibitor FLI-O6 (Krämer et al., 2013) and the GSIs LY-450139 (Semagacestat), LY-411573 and DAPT were tested in three doses (1, 5, 10 µM) resulting in a dose-dependent reduction of reporter signals (Supplemental Figure 2A). LY-411573 and DAPT reduced the signal most efficiently at 10 µM (46% and 49%) and LY-450139 and FLI-06 reduced the signal to 52% and 62% of DMSO (Supplemental Figure 2A), confirming DAPT as a reliable control to define the hit window; validation of normalization across plates and signal-to-background ratios, see Supplemental Figure 2B-F).

Screening of the 37,966 compounds identified 189 potential hits that matched the criteria: >70% reduction compared to DMSO; <25% difference from the Renilla-luc reporter (Figure 2A, Supplemental Table 1). The 189 compounds were subsequently tested in a three-dose regimen (1,25, 10 and 25 µM), along with the established Notch inhibitors and three luciferase inhibitors (PTC-124, luciferase inhibitor 1 and Nano-Luc inhibitor) as controls. As expected, the luciferase inhibitors showed the strongest quenching of the signal: the signal was reduced by 83% and 80% by PTC-124 and luciferase inhibitor 1, respectively. No signal reduction was observed for the Nano-luciferase specific inhibitor (Nano-Luc), underscoring the specificity of the inhibitors (Supplemental Figure 2G; for DMSO/DAPT signals and plate variations, see Supplemental Figure 2H-L). In the three-dose assay, 103 of the 189 compounds showed inhibition of reporter activity at 10 µM (which was defined as the cut-off), while 37 and 136 compounds showed reduced reporter activity at 1,25 and 25 µM, respectively (Figure 2B; Supplemental Table 1). In sum, after the initial screening of 37.966 compounds, 103 compounds matching the preset hit criteria remained.

**Figure 2:**
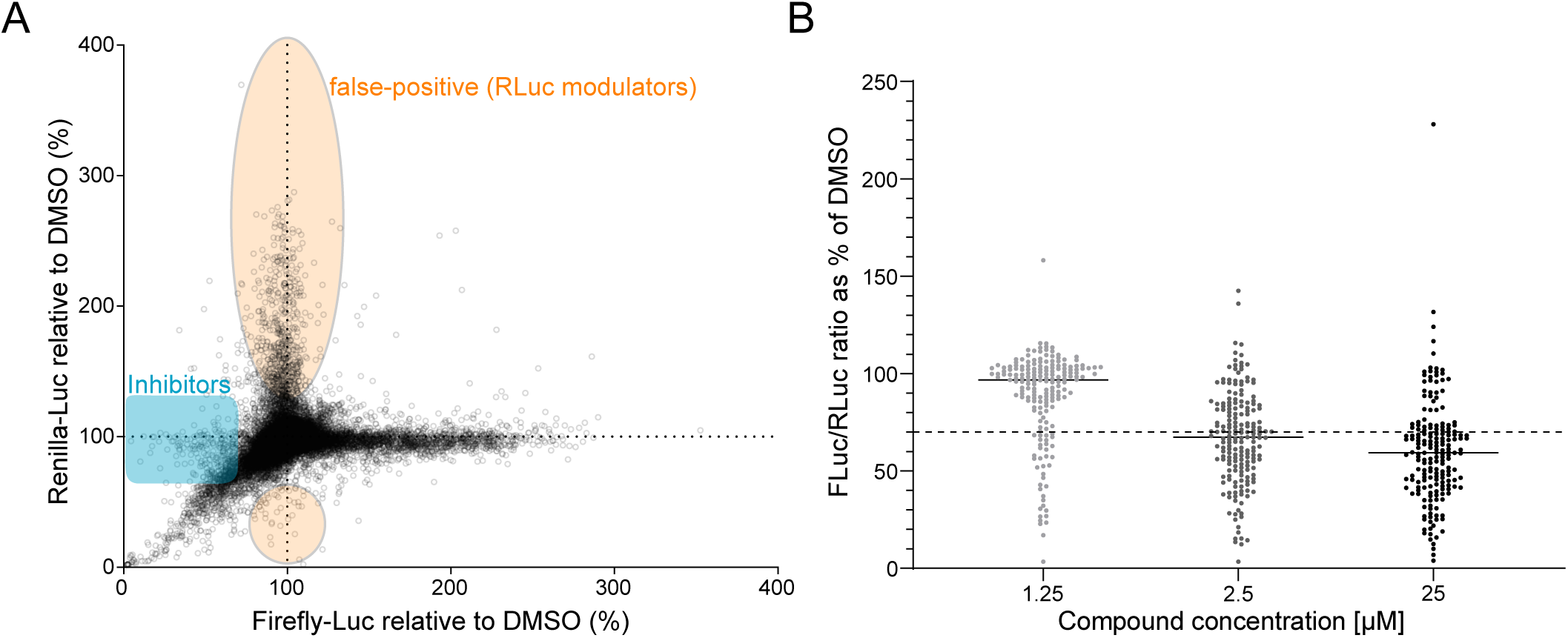
A 37,966 small compound library screen and compound confirmation assay. **(A)** A diagram showing the distribution of potential Notch inhibitors (decreased firefly luciferase activity; N=189; blue sphere) and false-positive compounds (with an increase also in the renilla luciferase activity; N=575; peach sphere). Firefly and renilla luciferase signals in % of DMSO are depicted on the x-axis and the y-axis, respectively. Screening was conducted after one day of compound exposure. The threshold for a Notch inhibitor was set to a decrease of firefly luciferase activity by more than 30% and no more than 25% increase in the activity of renilla luciferase. **(B)** Three dosage (1.25 10 and 25 mM) confirmation screen for the 189 inhibitors. The dotted line indicates the 70% reduction compared to DMSO cut off and 103 inhibitors were confirmed at 10 µM.

### Removal of potential γ-secretase inhibitors through an APP-based counter screen

As the initial set of hits might include potential GSIs, we performed a counter screen to remove such compounds, because GSIs are already well-established as pre-clinical pan-Notch inhibitors and have shown side effects in clinical trials (Andersson & Lendahl, 2014; Braune & Lendahl, 2016). To this end, we used a previously established reporter gene-based GSI assay (APP-C99) (Figure 3A), which measures γ-secretase activity through cleavage of amyloid precursor protein (APP), another protein processed by the γ-secretase complex (Karlström et al., 2002) (for analysis of assay stability and variation, effects of established Notch and luciferase inhibitors, see Supplementary Figure 3A,B). Using 70% signal reduction compared to baseline (DMSO) at 10 µM as threshold for removing a compound, 50 of the 103 compounds were removed as potential GSIs (Figure 3B, Supplemental Figure 3C, Supplemental Table 2).

**Figure 3:**
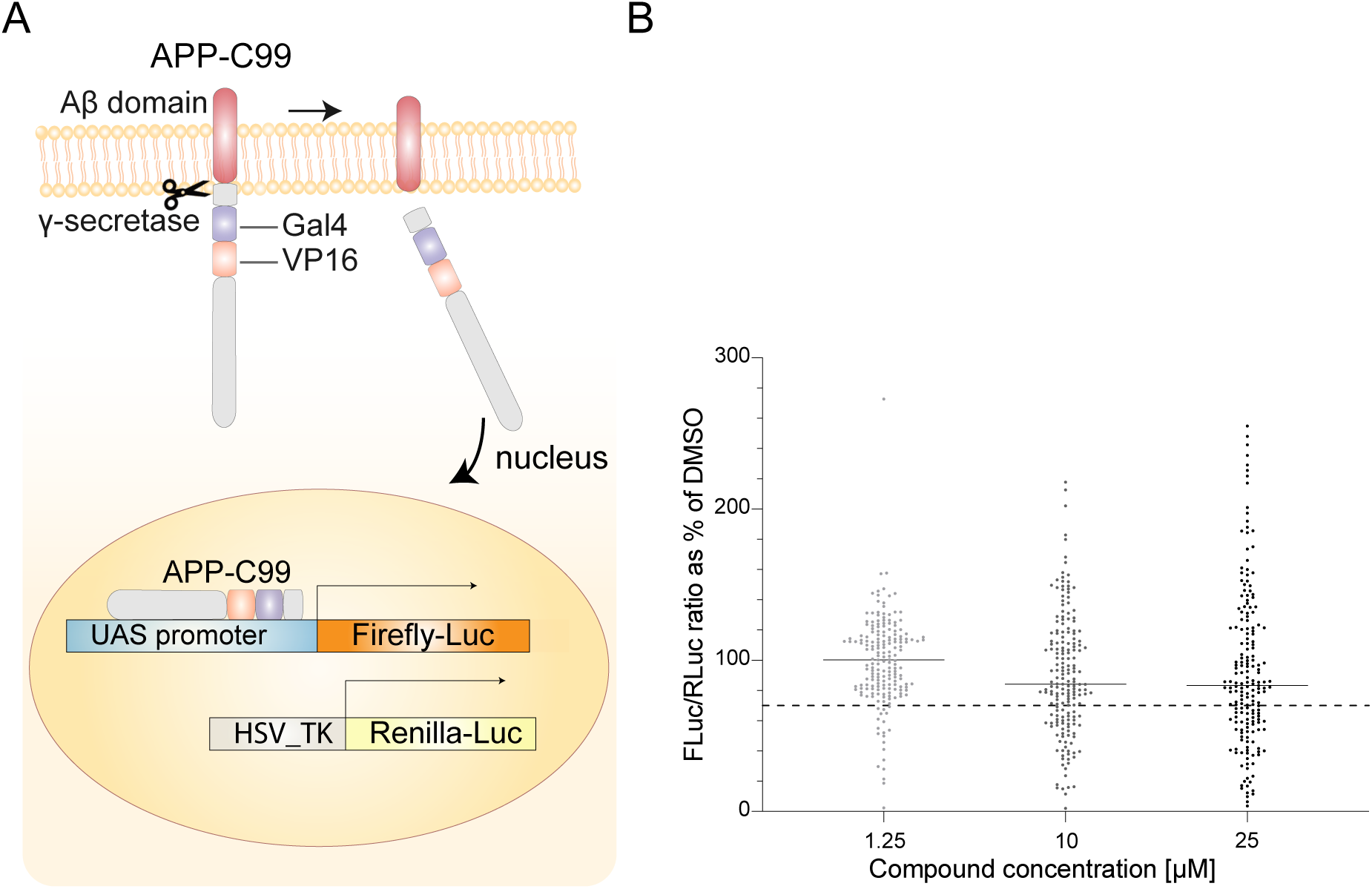
Analysis of Notch inhibitor candidates by the APP-C99 counter screen to eliminate potential GSIs. **(A)** Schematic depiction of the APP-C99 counter assay (Karlström et al., 2002). The engineered APP protein contains a GAL4/VP16 activation domain, which activates a UAS-firefly luciferase reporter gene construct upon cleavage of APP by the g-secretase complex. The HSV-TK promoter-renilla luciferase construct was used as a control for luciferase activity. **(B)** The 103 potential inhibitor hits from Figure 2B were passed through the APP-C99 counter screen at three dosages (1.25, 10 and 25 mM). The dotted line indicates the 70% reduction compared to DMSO cutoff and 53 potential inhibitor hits passed the APP-C99 counter screen at 10 µM.

The remaining 53 compounds were then filtered for undesired chemical properties, such as pan-assay interference compounds (PAINs), Rapid Elimination Of Swill (REOS, for review see (Walters & Namchuk, 2003) and compounds that generated false positive results and may act on luciferase directly were also removed. Together, 11 compounds were eliminated from further analysis, and the 42 remaining compounds were then subjected to an extended 11-point dose-response regimen (39 nM-40 µM) in the APP-C99 counter screen, leading to removal of an additional three compounds. Thus, 39 potential inhibitor hits remained for subsequent analysis (Supplemental Table 3; for inhibitor controls and dose response curves see Supplemental Figure 3D, E and for plate statistics see Supplemental Figure 3F, G).

### Identification of five inhibitor hits

To further narrow down the number of compounds, the 39 inhibitor hits passing the APP-C99 counter assay were scrutinized for various properties. Sixteen compounds predicted to have favourable pharmacogenetic, pharmacodynamic and toxicology properties based on their chemical structures were selected for subsequent experiments. These compounds were grouped based on their diverse core chemistries, and analogues from existing libraries were identified, resulting in five lead series. The selected analogues added 47 structurally related compounds to the 16 hits (for full list and chemical structures of the 16 compounds and respective analogues see Supplemental Table 4). All 63 compounds were analysed in the APP-C99 counter assay in an optimized 14-point dose-response analysis format, which led to removal of 11 compounds (one original compound and 10 analogues) that decreased the signal less than 70% compared to DMSO controls, leaving 52 compounds for subsequent analyses (for assay optimisation see Supplemental Figure 4A, for plate statistics and inhibitor controls see Supplemental Figure 4B and dose-response curves are shown in Supplemental Figure 4C).

Next, we subjected the 52 compounds to an optimized 14-point dose-response Notch reporter assay analysis, to establish inhibitory concentration 50 (IC50) values (for optimization of the assay using previously identified GSIs see Supplemental Figure 4D; for technical details of the analysis see Supplemental Figure 4E,F). Based on a 11-point pre-screen (data not shown), the 52 compounds were sorted into three baskets, each corresponding to an adjusted dose-range aiming to improve the curve fits and IC50s for the individual compounds in each basket. Of the 52 compounds, 22 showed an apparent signal reduction (14 of the original compounds and eight analogues, see Supplemental Table 5a for IC50 values). Five of the original 16 hit compounds, hereafter referred to as compounds A-E, exhibited IC50 values between 0.73 µM (compound E) and 4 μM (compound A) and Hill slopes between -16.2 (compound C) and -1,7 (compound D, Figure 4A, Supplemental Figure 4G); the structures of compounds A-E are shown in Figure 4B. Three analogues to compound B showed an IC50 < 2 µM while other analogues showed IC50s >4 µM or did not result in proper curve fits (Figure 4C). In sum, compounds A-E were selected for further analysis based on their structural properties and IC50 values.

**Figure 4:**
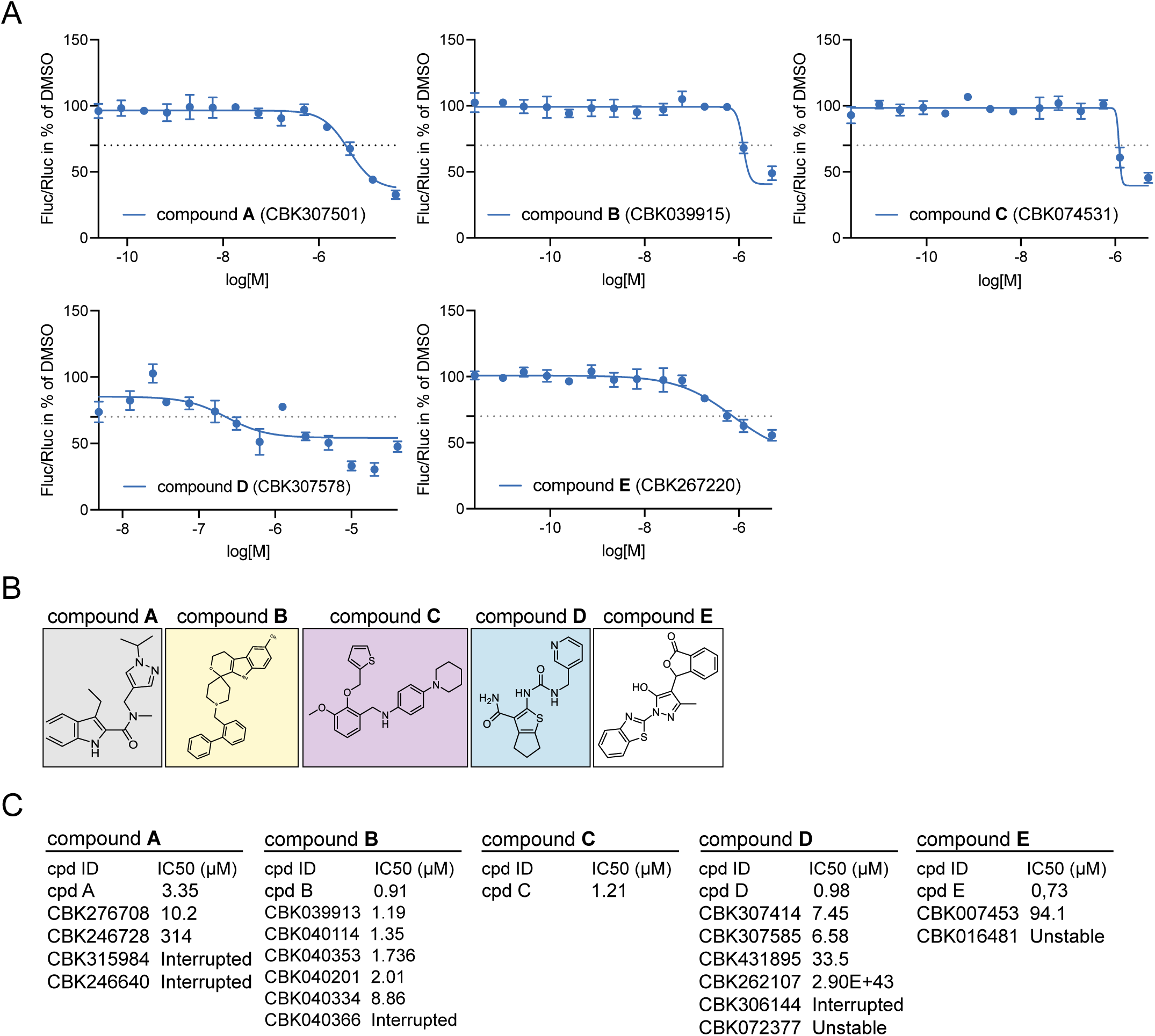
The five inhibitor hits A-E. **(A)** Dose-response curves from the Notch reporter assay using compound A-E at 14 different concentrations. **(B)** The chemical structures of compound A-E. **(C)** The IC50 values for the A-E compounds and names and IC50 values for analogues to the respective compound.

### Assessing Notch inhibitor hits A-E in an orthogonal assay based on myogenic differentiation

To corroborate their status as Notch inhibitors, we tested four of the five compounds (A, C, D, E) in an orthogonal assay for Notch function. Notch signalling plays an important role in myogenic differentiation (Vargas-Franco et al., 2022), and blocking Notch signalling promotes myogenic differentiation in the myoblast cell line C2C12 (Dahlqvist et al., 2003) (Figure 5A). C2C12 cells can be maintained as myoblasts but upon serum starvation spontaneously differentiate to MYH4-positive multinucleated myotubes, a process which is accelerated by Notch blockade (Dahlqvist et al., 2003). When subjected to the four compounds or to RO4929097 as GSI control for five days, C2C12 myoblasts showed elevated expression of the differentiation marker MYH4 for three of the candidate compounds (A, C, E) as well as for RO492097, accompanied by a transition towards a polynucleated myotube morphology (Figure 5B). Treatment with compound D did not produce elevated MYH4 levels but resulted in cell shape changes, with a transition towards a spindle-like morphology, indicating an onset of differentiation. The Notch inhibitor CB-103 (Lehal et al., 2020) produced increased MYH4 levels, but not reaching statistical significance (Figure 5B,C). Together, these data suggest that compounds A, C and E are endowed with Notch-inhibitory activity in the myogenic differentiation assay.

**Figure 5:**
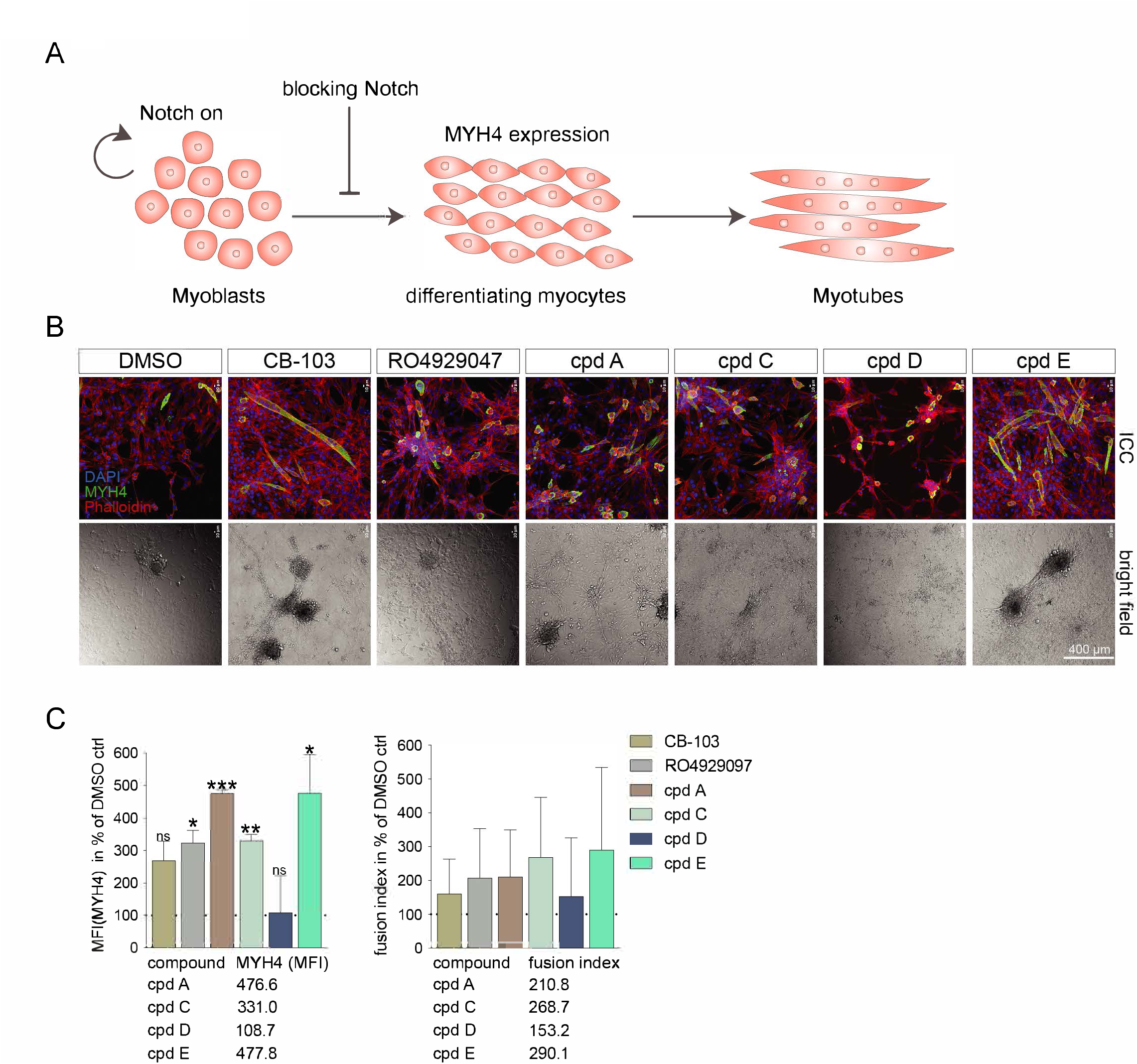
Analysis of compounds A-E in an orthogonal screen based on myogenic differentiation of C2C12 cells. **(A)** Schematic representation of the C2C12 myogenic assay system and how Notch blocks myogenic differentiation. **(B)** Immunocytochemistry (ICC) analysis of C2C12 cells five days after treatment with 10 mM of compound A, C, D and E, using DMSO as a reference and CB-103 and RO4929047 as controls (MYH4 antibody staining in green, Phalloidin in red, DAPI in blue). Below, bright field microscopy images of cells treated with compound A-E are shown. **(C)** Median fluorescence intensity of MYH4 (a myocyte marker) ICC staining in the C2C12 cells five days after treatment with 10 mM of compound A, C, D and E, and with DMSO as a reference. Treatment with CB-103 and ROI4929047 was included as controls. **(D)** Fusion of nuclei (as an indicator of differentiation) in the C2C12 cells five days after treatment with 10 mM of compound A, C, D and E, and with DMSO as a reference and CB-103 and RO4929047 as controls. *p ≤ 0.05, **p ≤ 0.01, ***p ≤ 0.001; ns = not significant.

### Analysis of growth-inhibitory effects in tumour cell lines

We next investigated whether compounds A-E affected growth and viability of cell lines from T-cell acute lymphoblastic leukemia (T-ALL) and breast cancer, two tumour types in which elevated Notch signalling has been implicated in tumorigenesis (Aster et al., 2017). We analysed the metabolic activity (CellTiter-Glo, Promega) in three T-ALL and six breast cancer cell lines exposed to 14 different concentrations (50 µM to 2.5 nM) of compounds A-E. Growth of the NOTCH1-addicted T-ALL cell lines JURKAT and RPMI-8402 was significantly reduced upon compound B or C treatment, resulting in IC50s of 274 nM and 7.12 µM in JURKAT cells and 614 nM and 713 nM in RPMI8402 cells after six days, for compound B and C, respectively (Figure 6A). Furthermore, the NOTCH3-addicted TALL-1 cell line showed growth inhibition after treatment with compound B for six days with an IC50 of 7.2 µM. As control, the Notch inhibitor CB-103 reduced growth with similar potency as compound B and C in JURKAT and RPMI-8402 cells (IC50 1.78 µM and 927 nM after six days) and reduced growth of TALL-1 cells most efficiently after 6 days (IC50 290 nM). When testing analogues of compounds B and C, five of the seven B analogues and one of the two C analogues produced dose-response curves and IC50s, which however were higher compared to the original hits (Supplemental Figure 5A; IC50s are presented in Supplemental Table 5b).

**Figure 6:**
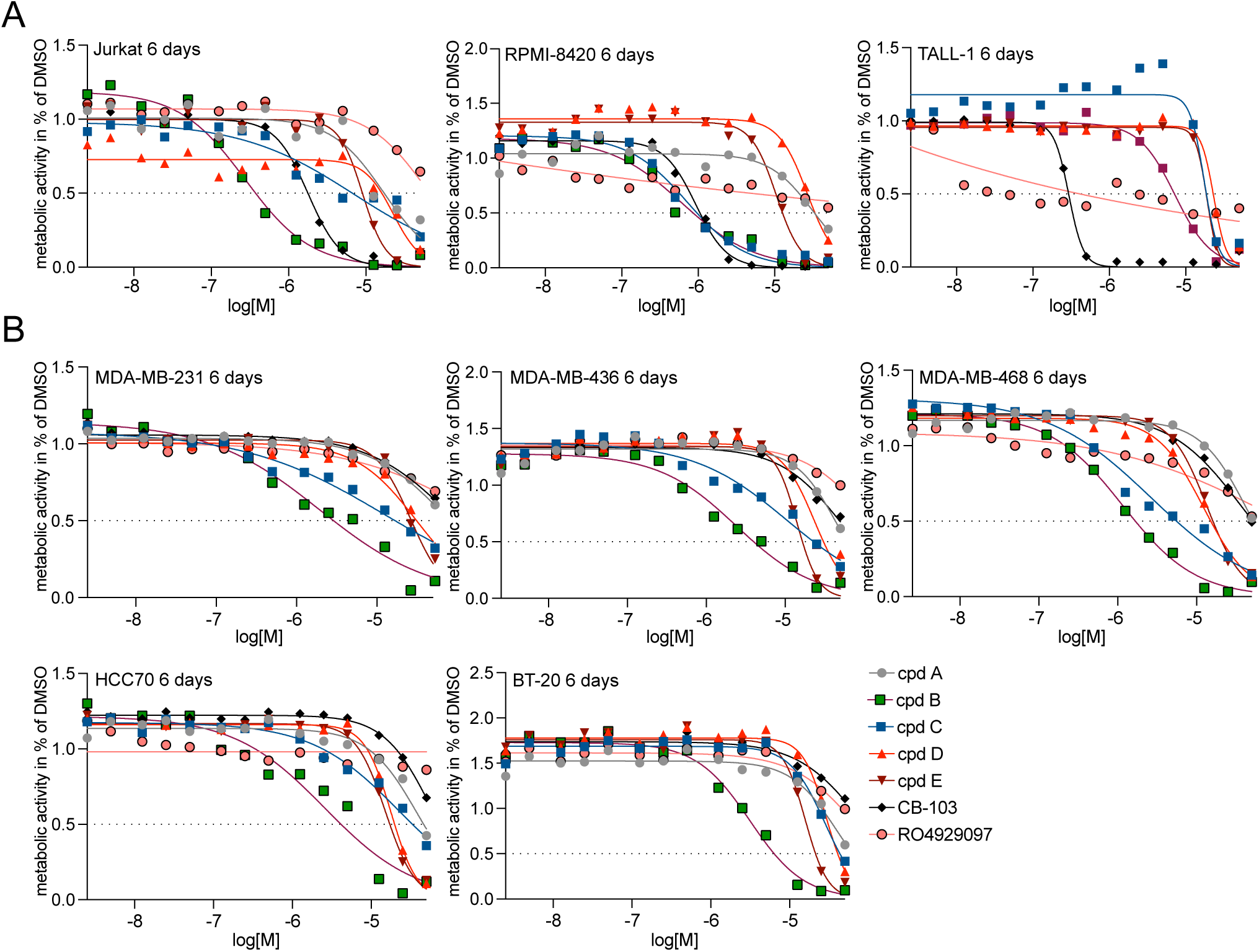
Analysis of compounds A-E for growth-inhibitory effects on T-ALL and breast cancer cell lines. **(A)** Graphs showing 14-point growth-response curves for the level of metabolic activity in three T-ALL cell lines (JURKAT, RMPI-8420 and TALL-1) at six days after treatment with compound A-E, and CB-103 and RO4929097 as controls. **(B)** Graphs showing 14-point growth-response curves for the level of metabolic activity in five breast cancer cell lines at 6 days after treatment with compound A-E and CB-103 and RO4929097 as controls. Results of two biological assays were averaged for potential hit compounds.

Growth of the breast cancer cell lines MDA-MB-231 and HCC70 was only marginally affected by compound A-E treatment after three days (IC50s >25 µM) (Supplemental Figure 5B, Supplemental Table 5b), but after six days of treatment with compound B, there was a robust decrease in metabolic activity in the MDA-MB-231, MDA-MB-468, MDA-MB-468, HCC70 and BT-20 cells (IC50s of 1,9 µM, 1,2 µM, 2,4 µM, 2,6 µM and 3,1 µM, respectively). Treatment with compound C for six days resulted in an IC50 of 2.7 µM in MDA-MB-468 cells, but higher IC50s (>10 µM) in the other breast cancer cell lines (Figure 6B). Compound A, D and E treatment resulted in IC50 values >10 µM for all cell lines. We also tested three analogues to compound B and one analogue to compound C. The compound C analogue produced a dose-response curve fit with IC50 > 5 µM in MDA-MB-468 cells and > 10 µM in MDA-MB-231 cells, whereas the compound C analogue produced IC50s >10 µM in MDA-MB-468 and MDA-MB-231 cells (Supplemental Figure 5C; for IC50s see Supplemental Table 5b).

Next, we were interested in learning whether compounds A-E affect expression of Notch downstream genes. Analysis of Notch downstream gene (HES1, c-MYC, JAGGED1 and HES4) expression in the tumour cell lines revealed a dose-dependent downregulation of HES1 and c-MYC expression in T-ALL cells upon treatment with compound B, C, D and E (Supplemental Figure 6A). HES1 was downregulated in MDA-MB-231 cells when treated with compound A, D and E, and expression of HES4 was reduced after treatment with A-D, while the transcript levels of the Notch ligand Jagged1 were reduced when cells were treated with compound A, B, D and E in MDA-MB-231 cells. CB-103 treatment unexpectedly led to upregulation of c-MYC in the MDA-MB-231 cell line and upregulation of HES1 and HES4 in MDA-MB-468. In contrast, the GSI RO4929097 consistently reduced target gene expression across the panel of cell lines but led to upregulation of c-MYC expression in MDA-MB-231 cells (Supplemental Figure 6B,C). In sum, compounds B and C induced growth inhibition and downregulation of Notch target genes in several breast cancer and T-ALL cell lines.

### Compound C is a dihydroorotate dehydrogenase inhibitor

Compound C had an overall attractive profile, accelerated myogenic differentiation and potently inhibited tumour cell line growth, and we therefore decided to explore it in more detail. When comparing its structure to that of other known classes of inhibitors, we noted that compound C was partially structurally related to an earlier-generation inhibitor of dihydroorotate dehydrogenase (DHODH) (Leban et al., 2004; Lolli et al., 2012) (Figure 7A). DHODH is an enzyme localized to the inner mitochondrial membrane, where it converts dihydroorotate to orotate in the de novo pyrimidine synthesis pathway (Madak et al., 2019). The fact that a genetic link between Notch and DHODH signaling had previously been published (Thörig et al., 1987) further spurred us to test whether compound C was endowed with DHODH-inhibiting properties. In keeping with this notion, compound C reduced the reporter assay activity at 1 and 10 μM concentrations, and this effect could be reversed by addition of 100 μM uridine (Figure 7B), which is known to abrogate the effect of DHODH inhibitors (Ladds et al., 2018). Furthermore, a well-established DHODH inhibitor, BAY2402234, reduced the activity of the Notch reporter activity at 0.1 μM, an effect that could also be blocked by addition of 100 μM uridine (Figure 7B). These experiments demonstrate that compound C acts on Notch signalling via its DHODH inhibitor function and that another DHODH inhibitor also can block Notch signalling.

**Figure 7:**
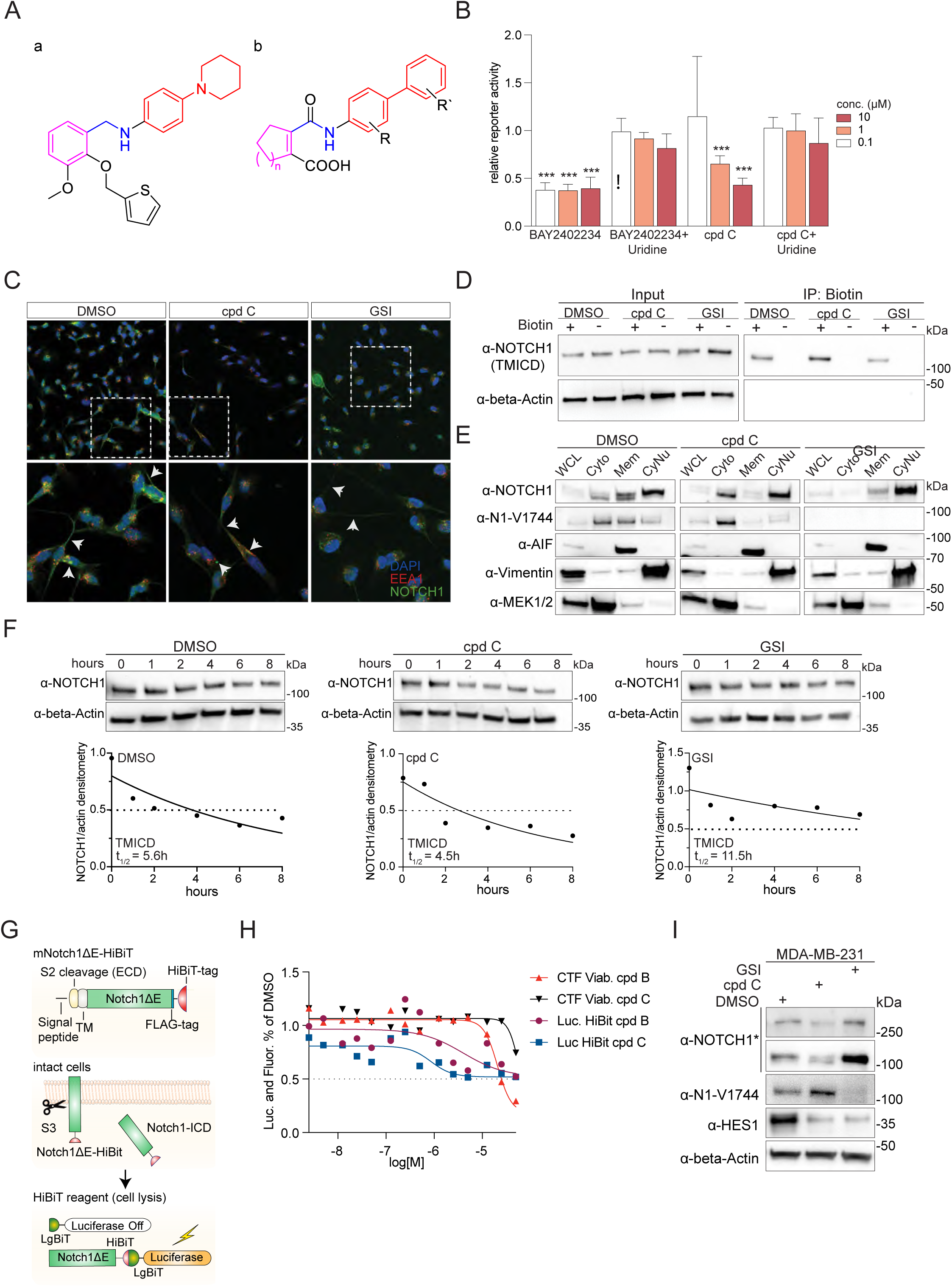
Compound C represents a DHODH inhibitor. **(A)** Chemical structures of compound C and a previously proposed potent DHODH inhibitor (Lolli et al., 2012). The colors indicate similarities between the compounds, with two hexacyclic ring arrangements (red) connected through an amine-like linker (blue) to a second ring arrangement (magenta). **(B)** The levels of Notch reporter activity after 24h treatment with DHODH inhibitor BAY2402234 and compound C at 0.1, 1 and 10 µM in presence or absence of 100 mM Uridine. Bars represent reporter activity relative to DMSO as control for the BAY2402234 and the cpd C columns and with DMSO+100 mM Uridine as control for the BAY2402234+Uridine and the cpd C+Uridine columns. Bars are presented as mean and error bars display SD of three biological assays. Changes were considered significant with *p<0.05, **p<0.01, ***p<0.001. **(C)** The localization of Notch1 was determined by immunocytochemistry in MDA-MB-231 cells, using an antibody (NOTCH1, in green) recognizing both the full length, TMICD and ICD forms of NOTCH1. The localization in response to treatment with 5 µM of compounds B and C or with a GSI (RO4929097) for 24 hours, as indicated, is shown. Early endosomes were visualized by an antibody to EEA1 (in red), and nuclei were stained by DAPI (in blue). The boxed areas are shown as high magnifications in the lower row. **(D)** Western Blot analysis of surface biotinylation experiment of MDA-MB-231 cells after 24h treatment with compound C or GSI, as control. Cells were either subjected to biotinylation or left untreated as controls (input, left panel), followed by pull-down of biotinylated proteins (right panel). Results were visualised with anti-NOTCH1 antibody, and beta-actin antibody as loading control. **(E)** Western blot analysis of intracellular distribution of TMICD and ICD following subcellular fractionation (cytoplasmic (Cyto), membrane plus organelles (Mem), cytoskeletal and nuclear and (CyNuc), as well as whole cell lysate (WCL) from MDA-MB-231 cell extracts. Cells were treated with 2 µM of compound C or 2 µM of GSI (RO4929097) for 24 hours. The Western blot was stained with an antibody recognizing both the full length, TMICD and ICD forms of NOTCH1 (a-NOTCH1; upper) or with an antibody recognizing the neoepitope appearing only at the S3 cleaved Notch1 ICD (a-N1-V1744) (lower). Mitogen-activated protein kinase kinase 1/2 (MEK1/2), Apoptosis-inducing factor (AIF) and Vimentin (Vim) antibodies were used as controls to label the respective cell fractions (Cyto, Mem, and CyNuc, respectively). **(F)** Analysis of cleaved NOTCH1 longevity by Western blot in the 293T Notch reporter cell line (#A1-11) 0-8 hours after addition of cycloheximide (CHX) to block translation. The effects of DMSO and 5 µM compound C and GSI (RO4929097) are shown. A pan-NOTCH1 antibody was used to visualise NOTCH1 bands and beta-actin staining was included as loading control. Graphs at the bottom show calculation of half-lives for cleaved NOTCH1 after CHX treatment from Western blot. The half-life reduction is expressed relative to NOTCH1 in DMSO control. **(G)** Schematic depiction of the HiBiT assay. The Notch1DE-HiBiT construct is depicted on top, and the cleavage by g-secretase is shown in the middle panel. At the bottom, the interaction with the LgBiT moiety, generating luciferase activity, is shown. **(H)** Graphs showing 14-point growth-response curves for the level of HiBiT luciferase activity in the HiBiT reporter cell line of two averaged experiments 24h after treatment with compound B and C at dose-ranges from 50 mM to 25 nM. In addition, dose-response curves for cell viability (labelled CTF Viab; using the CellTiter-Fluor Viability assay, CTF) are shown. **(I)** Western blot analysis of NOTCH1 and HES1 protein levels in MDA-MB-231 in response to 48h treatment with DMSO, 2 µM compound C and GSI (RO4929097). Beta-actin staining was included as loading control. The a-NOTCH1 antibody recognizes both the full length, TMIC and ICD forms of NOTCH1, with the full-length form at approximately 250 kDa and processed forms at around 100 kDa. Activated NOTCH1 was visualized using the antibody targeting the S3 cleaved form of NOTCH1 (a-N1-V1744). The asterisk denotes that the Western blot membrane was stripped and reprobed with the a-NOTCH1 antibody a-N1-V1744.

After demonstrating that compound C is a DHODH inhibitor, we explored how compound C inhibited the Notch signalling pathway. In compound C-treated MDA-MB-231 cells, NOTCH1 was detected at the cell surface as well as in cytosol and nucleus, indicating that subcellular distribution is not overtly affected (Figure 7C). In contrast, treatment with 5 µM GSI (RO4929097) resulted in increased NOTCH1 membrane localisation (Figure 7C), in keeping with the role of γ-secretase in NOTCH1 processing and release from the cell membrane. The analysis of intracellular localization was complemented by cell surface biotinylation experiments supporting the notion that NOTCH1 surface presentation is not hindered in compound C-treated cells (Figure 7D). To corroborate the intracellular localization, we analysed the distribution of the TMICD and ICD forms of NOTCH1 in MDA-MB-231 cells by cell fractionation experiments (Figure 7E, see point 1b above). NOTCH1 localised to the cytosolic, membrane and nuclear fraction in control samples, while predominantly localised to the cytosolic and the nuclear fraction in compound C-treated cells. The distribution of S3-cleaved Notch1 ICD, detected by an antibody specifically recognizing the neoepitope (Val 1744) after S3-cleavage (Chapman et al., 2006), differed between DMSO and compound C-treated samples: compared to the DMSO control samples, the ICD was more abundant in the cytosolic fraction, while only sparsely detectable in the membrane fraction, following treatment with compound C. No ICD bands were detected in GSI treated samples that was used as another control, as expected (Figure 7E).

We also asked whether the stability of the cleaved form of the receptor was affected by compound C. Cycloheximide experiment to measure the half-life of NOTCH1 showed that the cleaved form of the receptor is more stable in DMSO controls compared to compound C treated cells (t_½_ 5.6h, respectively 4.5h) (Figure 7F). Treatment with a GSI resulted in enhanced protein stability (Figure 7F), which is in line with the expected accumulation of unprocessed protein within the cells. To provide evidence that compound C may reduce Notch1 levels, we assessed the amount of Notch1 ICD in an alternative manner, using a mouse Notch1ΔE (mNotch1ΔE) moiety (Chapman et al., 2006), which spontaneously generates mNotch1 ICD. After engineering a stable cell 293T cell line with mNotch1ΔE fused to an HiBiT tag (Figure 7G), exposure to compound C in a 14-point dose-response assay resulted in a dose-dependent decrease of luciferase signal (Figure 7H, Supplemental Table 5b); for signal stability and variation see Supplemental Figure 7A; for corroborating data from a second clonal cell, see Supplemental Figure 7B). As controls, treatment with Bafilomycin A1, which prevents protein degradation by inhibiting autophagosome-lysosome fusion (Yamamoto et al., 1998) or the proteasome inhibitor MG132 increased the HiBiT signal (Supplemental Figure 7C). To determine whether the reduced levels of Notch1 ICD resulted in lowered levels of Notch downstream expression, we assessed the amount of HES1 in MDA-MB-231 cells, and treatment with 2µM compound C for 48h caused a strong reduction in HES1 levels (Figure 7I). The total amount of NOTCH1 receptor (both the full length 250 kD and the TMICD 100 kD form) appeared to be somewhat reduced after compound C treatment in the MDA-MD-231 cells (Figure 7I). In conclusion, these data suggest that compound C does not hinder surface presentation or processing of the NOTCH1 receptor but may reduce protein half-life.

## Discussion

Hyperactivated Notch signalling is linked to several types of disease and cancer, and there is thus a need to develop novel Notch inhibitors (Andersson & Lendahl, 2014; Majumder et al., 2021). In this report, we conducted an unbiased large-scale small molecule screen to identify potential Notch inhibitors based on a novel reporter assay for immediate downstream Notch signalling as the read out. An initial screening of 37.966 compounds was followed by a counter screen for GSI activity and an orthogonal screen for blockade of myogenic differentiation, which eventually led to the identification of five hits (compounds A-E); the stepwise screening is schematically depicted in Figure 8. The five hits have distinct chemical structures, indicating that they may target Notch signalling by distinct mechanisms and that the screen indeed can capture different types of Notch inhibitor candidates.

**Figure 8:**
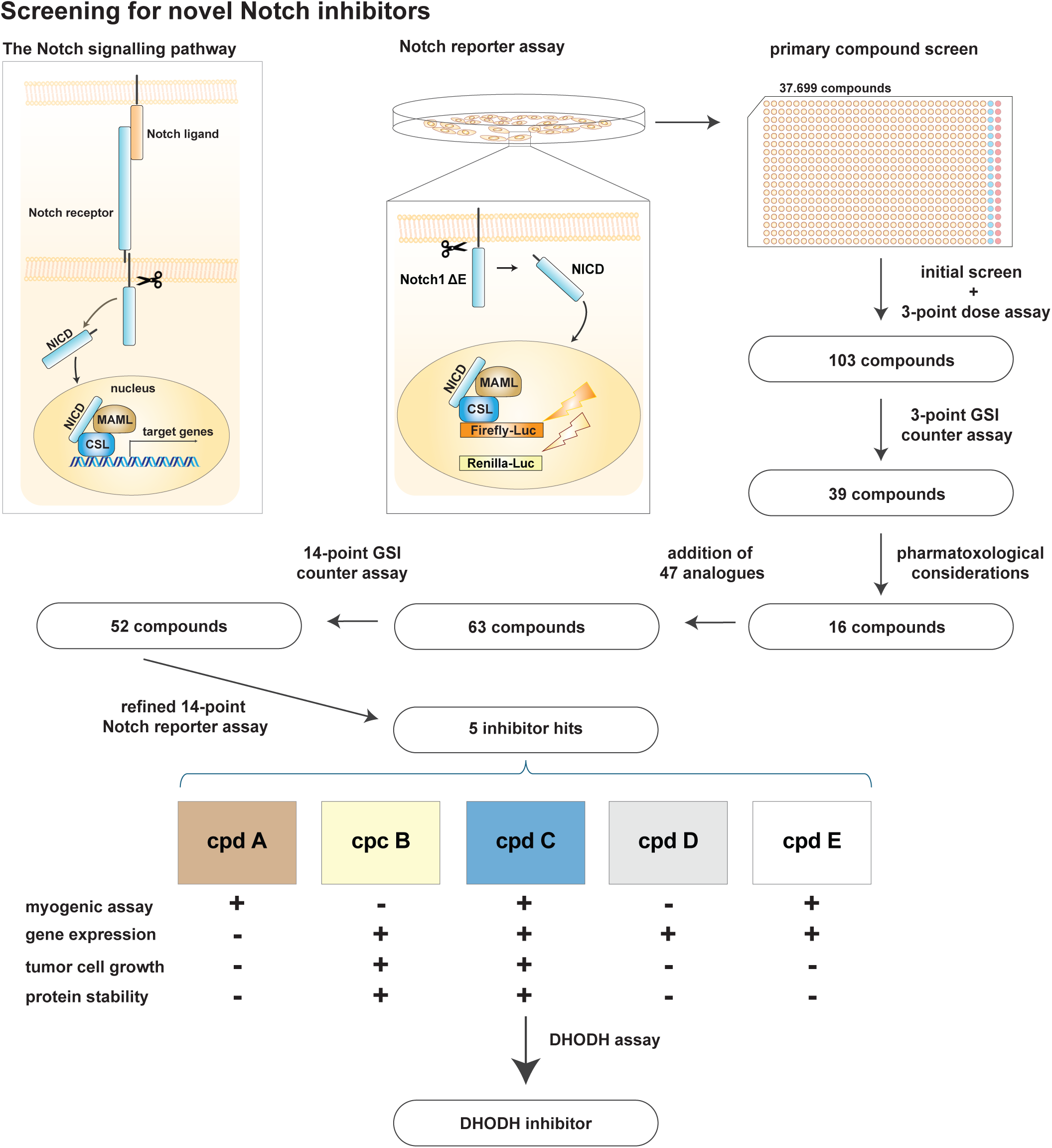
A summary of the small compound screening strategy leading to the identification of compound C as a novel DHODH inhibitor. In the upper panel the Notch signalling pathway (left) and the reporter assay (middle) are shown. To the right in the upper panel and in the middle panel the various steps in the compound screening procedure are depicted. The numbers in the boxes illustrate the number of candidate hits after each screening step. In the bottom panel, the structures of the five selected compounds (compounds A-E) are shown, and below, their performance in the various deconvolution steps, as indicated (+ indicates that the compound had an effect; - indicates that there was no significant effect in the respective assay). When assaying for DHODH inhibitor activity, compound C turned out to be a novel DHODH inhibitor.

Among the five inhibitor hits, we focused our attention on compound C, which showed pronounced structural similarities to dihydroorotate dehydrogenase (DHODH) inhibitors. Confirmation of compound C as a DHODH inhibitor was achieved by uridine treatment which abrogated the inhibition of Notch signalling by compound C in the Notch reporter assay, in line with DHODH’s critical role for pyrimidine production (Madak et al., 2019). Furthermore, a well-established DHODH inhibitor, BAY2402234, reduced Notch signalling in the reporter assay, and collectively these data underscore that compound C exerts its effect on Notch via its DHODH inhibitor function. This link between DHODH and Notch signalling is consistent with several earlier studies. In the fruit fly, *Drosophila melanogaster*, chemical DHODH inhibition by 5-methyl orotate phenocopied *Notch* mutations, including notches of wings and bristle multiplication (Thörig et al., 1987). Furthermore, it was recently shown that DHODH inhibition, via administration of the DHODH inhibitor brequinar, was curative in a Notch1 ICD-induced mouse model of T-ALL (Sexauer et al., 2023). Finally, a recent report demonstrated that increased expression of DHODH may lead to elevated levels of Notch1 ICD (He et al., 2022). Together, these studies substantiate a link between DHODH function and Notch signalling and supports the notion that DHODH inhibition may be an interesting avenue to explore for novel Notch inhibitors.

The mechanistic coupling between DHODH function and Notch signalling largely remains to be elucidated, but our data from deconvolution studies of compound C reveal a reduction in the amount of NOTCH1 ICD in MDA-MB-231 cells, which may indicate a decreased stability and more rapid turnover of NOTCH1 ICD. Alternatively, compound C may cause a shift in localization, although the overall subcellular localization of Notch receptors is however likely not affected, as the distribution of NOTCH1 between the cell surface, the cytosol and the nucleus was not significantly altered, arguing against differences in receptor processing rates. As the DHODH enzyme is primarily localized to the inner mitochondrial membrane it is unlikely that it directly affects Notch receptor stability. The reduction of NOTCH1 ICD levels may therefore be an indirect effect of DHODH acting via proteins that posttranslationally modify Notch ICD and reduce Notch ICD stability, for example kinases such as PIM kinases (Landor et al., 2021; Santio et al., 2016) and E3 ubiquitin ligases such as cdc4 (Oberg et al., 2001). Hydroxylases or sumoylases may also be involved as intermediates (Antfolk et al., 2019). Another possibility is that the effect on Notch instead is mediated by downstream effects of DHODH inhibitors. For example, it has been demonstrated that DHODH inhibitors block the degradation of p53 (Ladds et al., 2018), and there is evidence for a functional relationship between Notch signalling and p53 (Jin et al., 2012; Yun et al., 2015). Furthermore, DHODH inhibitors may regulate MYC (Tsea et al., 2024), which also is linked to Notch signalling (Chan et al., 2007; Palomero et al., 2006), and a downregulation of c-MYC is observed after compound C treatment in MDA-MB-231 cells. Recently, it has also been shown that DHODH inhibitors induce specific metabolomic changes in lung cancer cell lines (Schuhknecht et al., 2025), but whether this affects Notch signalling remains to be investigated. Alternatively, as addition of uridine reversed the effect of compound C, it is also possible that the role of DHODH in pyrimidine nucleotide synthesis is somehow related to the effect of compound C on Notch signalling. Additional research will therefore be required to decipher the precise mechanistic link between Notch and DHODH.

The finding that DHODH inhibitors reduce Notch signalling is also of importance for a better understanding of potential side effects emerging from off-target Notch inhibition when DHODH inhibitors are used in specific disease settings. DHODH inhibitors, such as leflunomide and teriflunomide, are currently used in therapy for rheumatoid arthritis, multiple sclerosis, psoriasis and diabetes (Madak et al., 2019; Reis et al., 2017; Zhang et al., 2021) and the range of diseases in which DHODH inhibition may be beneficial is increasing: there is an emerging interest in DHODH inhibition in malaria treatment (Chinnappanna et al., 2023; Nie et al., 2025) and for acute myeloblastic leukemia therapy, where DHODH inhibition may enhance myeloid differentiation (Sharma & Borthakur, 2023). In these situations, it is important to learn about potential consequences of the DHODH inhibitors simultaneously reducing Notch signalling; adverse effects of off-target reduction of Notch signalling have been painfully learned from the use of GSIs in clinical trials for Alzheimer’s disease. As GSIs not only block APP processing but also abrogate Notch receptor cleavage, patients in such trials showed reduced Notch signalling, accompanied with for example gastrointestinal toxicity, immune suppression and increased risk for skin cancer (Andersson & Lendahl, 2014).

In conclusion, we show that an unbiased small compound screen monitoring the level of Notch downstream activation yielded distinct types of inhibitor hits and importantly unravelled a link between DHODH and Notch signalling. This will lay the foundation for further exploration of novel Notch inhibiting strategies based on DHODH inhibition and may also inform on potential Notch off-target effects from the current use of DHODH inhibitors in the clinic.

## Materials and Methods

### Generation of the Notch reporter construct and the reporter cell line

A de novo Notch reporter construct, composed of six dimeric CSL binding sites linked to a basic β-globin promoter and followed by a firefly luciferase gene and a PEST degradation domain was synthesised (GeneArt). The construct was stably introduced into 293T cells, and zeocin (0.1 mg/ml) or Puromycin (0.05 mg/ml (ThermoFisher, cat #R25001, #A1113803) were used as selection markers. A clonal cell line (clone #A1) with robust renilla expression and enhanced firefly luciferase expression upon transient transfection of Notch1 ICD or Notch1ΔE constructs was selected, and a UB6 promoter-Notch1ΔE construct was stably introduced into this cell line using Blasticidin (0.1 mg/ml; ThermoFisher, cat #A1113903) as a selection marker. Stable cell lines were generated by serial dilution and only clonal colonies were selected for further evaluation. To measure luciferase reporter activity, cells were washed in cold PBS, detached, and collected in pre-warmed cell culture medium and centrifugated for five min at 200xg. The cell pellet was resuspended in medium, cells were counted, and 4.000 cells were seeded in each well of a white 384 well plate (Corning #3670) in 20 µl medium containing either DMSO or the GSI DAPT (N-[N-(3,5-Difluorophenacetyl)-L-alanyl]-S-phenylglycine t-butyl ester, Sigma-Aldrich, cat #565784-M) and incubated for 24h. The efficacy of the reporter gene system was determined by monitoring the level of luciferase activation in the absence or presence of different amounts of DAPT in several cell clones. Luciferase activity was assessed using the dual-GLO luciferase detection kit from Promega (cat # E2920), following the manufacturer’s instructions. A minimum of two technical replicates were measured per clone using a SpectraMaxi3 plate reader (Molecular Devices). Reporter activity of different clones and in response to DAPT was assessed by normalising firefly (Notch activity) to renilla luciferase signals. Normalised signals of individual clones cultivated in DMSO were then compared to corresponding signals generated with DAPT-treated cells. The cell clone with the strongest signal to control ratio (DMSO to DAPT) was then selected for further exploration (clone #A1-11).

### Generation of the HiBit reporter construct and cell line

The Notch1DE construct used to generate the Notch reporter cell line (see above) was first cloned in frame into the pBiT3.2-C [TK/HiBiT/Blast] Vector (Promega) and then stably introduced into 293T wild type cells, using Lipofectamine transfection, as described. Analogous to the Notch reporter cell line, Blasticidin was used as a selection marker (0.1mg/ml) and stable cell lines with robust luciferase signals were generated. HiBit signals correlated with cell numbers and clone #A4F2 showed the overall best luciferase signal to control and signal-to-noise ratio and was selected for further analysis. A clone (#D2) with an empty HiBit vector stably integrated was used as control.

### RNA isolation and qRT-PCR

Total RNA from cell lines was isolated using the RNeasy Mini kit (cat #74104, Qiagen), according to the manufacturer’s instructions. qRT-PCR was performed as described elsewhere (E.-B. Braune et al., 2016) with some minor modifications: RNA was reverse transcribed to cDNA using iScript Reverse Transcription Supermix (cat # 1708841, Bio-rad). cDNA was then diluted 1:5 in nuclease free H_2_0. qRT-PCR was performed on a C1000 Touch thermal cycler (Bio-rad) and gene expression was detected with SsoAdvanced Universal SYBR Green Supermix (cat # 1725274, Bio-rad). Gene expression was determined by normalizing to GAPDH mRNA expression levels and fold expression change was calculated using the ΔΔCT method (ΔΔCT=ΔCt sample-Δct control). Primers used in qPCR are shown in Supplemental Table 6.

### Cell lines

The breast cancer cell lines, HCC70, MDA-MB-468, and MDA-MB-231 and the T-ALL cell line Jukart were acquired from ATCC (cat #CRL-2315, #HTB-132, #CRM-HTB-26, #TIB-152), and the T-ALL cell lines RPMI-8402 and TALL-1 were acquired from DMSZ (cat #290, #521). Breast cancer and T-ALL cell lines were maintained in RPMI 1640 medium (ATCC’s modification, cat #A1049101, ThermoFisher) with 10% fetal bovine serum (FBS, heat inactivated, cat #A3840402, ThermoFisher) and 1% penicillin-streptomycin (cat #15140-122, Thermo Fisher Scientific). The 293T cell lines used in the small compound screening assays, including the Notch reporter cell lines and counter assay cell lines, were maintained in DMEM (ThermoFisher, cat #11995065) supplemented with 1% GlutaMAX (ThermoFisher cat #35050061), 10% fetal bovine serum (FBS, heat inactivated, cat #A3840402, Thermo Fisher) and 1% penicillin-streptomycin (cat #15140-122, Thermo Fisher). The 293T cell line used in orthogonal assay was cultivated without addition of GlutMAX. Accutase (cat # A6964-100ML, Sigma-Aldrich) was used to detach the cells prior to seeding on coated plates and during maintenance. All cell lines were cultivated with 5% CO_2_ at 37 degrees. C2C12 cells were obtained from Sigma-Aldrich (cat #91031101) and maintained in proliferation DMEM high glucose-based medium (ThermoFisher, cat #12430054) containing 20% FBS and 1% PenStrep (100U/ml) and were splitted 1:5 upon reaching 50% confluence. Cells were allowed to differentiate in DMEM high glucose medium containing 2% donor equine serum (HyClone cat #SH30074.02). Prior to seeding, cells were counted using automated cell counting devices. Scepter 2.0 (MerckMillipore) was used for counting cells for small compound screens and Bio-rad’s TC20 cell counter was used for cell counting in orthogonal assays.

### Transfection of cells and generation of clones

Cells were transfected using Lipofectamine 3000 (ThermoFisher, cat #L3000001), according to the manufacturer’s instructions. Two days post transfection, cells were detached, counted, diluted and single cells were seeded into 96-well plates. Seven days post seeding, wells containing monoclonal colonies were labelled and medium was changed. Clones were cultivated to 90% confluence and then further expanded.

### Screening of the small compound library

37,966 compounds were spotted by the compound center at SciLifeLab (https://www.scilifelab.se/) on a total of 120 plates (Greiner #781080) at a fixed concentration of 10 µM and the screening process spanned five consecutive sessions. The layout of each plate contained one column for DMSO and one for DAPT controls to determine the quality and reliability of the data. DAPT controls were spotted at 10 µM and DMSO was spotted in an equimolar volume. In addition, plates spotted with only DMSO were utilized to investigate the impact of plate effects before and after each session. Furthermore, an alternative set of controls, including several Notch inhibitors and GSIs, was used to verify the functionality of the assay and to ensure that its performance aligned with the expected dynamic downregulation upon blockage of Notch. Compound-spotted plates were stored at -20°C and were equilibrated to room temperature, followed by one minute centrifugation at 1000x g before seeding of cells. 293T Notch reporter cells were grown to 75% confluence, detached and 20 µl containing 4.000 cells were seeded in each well of a 384-well compound plate using automated cell dispenser systems (Multidrop, ThermoFisher). 24h prior seeding, dispenser cassettes were equilibrated, and tubing was thoroughly flushed with several washes of PBS followed by medium. Tubing was left in medium for 10 min before seeding cells to equilibrate conditions and ensure even dispensing. Cells seeded on compound plates were cultivated for 24h and placed in a box within the cell incubator with additional tissues soaked in sterile H_2_0 to avoid evaporation from compound plates. Luciferase reporter activity was assessed using the dual-GLO luciferase kit from Promega (cat #E2940). First, three compound plates per time were allowed to equilibrate to room temperature. Next, 20 µl firefly luciferase substrate was added to each well using an automated dispenser systems (Multidrop, ThermoFisher) followed by incubation with thorough agitation for 10 min in the dark. Firefly luciferase signals were then measured on a Wallac Victor3 1420 plate reader (Perkin Elmer) with a signal integration time of 0.5 seconds. After reading firefly signals of all compound plates, the dispenser cassette was changed to avoid contamination between the different reporter signals. Next, 20 µl of renilla firefly substrate was added as before and plates were incubated for 10 min in the dark while shaking. Subsequently plates were incubated for an additional 10 min in the dark before signals were read. Activity of compounds was assessed by calculating the ratio of firefly and renilla for each well, followed by normalising each value to the averaged DMSO control values of each plate. Normalised DMSO and DAPT control values were determined, and the respective standard deviation was then used to calculate the Z-factor for each plate to determine the quality of the screen and assay window. A Z-factor of >0.5 was considered excellent.

Inhibitor hits were selected by setting the threshold close to a theoretical exclusion criterium. The threshold was defined by first calculating the sum of the three-fold standard deviation (SD) for normalized DMSO respectively DAPT controls, averaged across all plates. The sum of the averaged inhibition by DAPT and the combined three-fold SDs of DAPT and DMSO was then calculated. This theoretical criterium threshold was 72%, meaning that compounds that do not reduce the signal to more than 72% of DMSO should be excluded. The hit threshold was set to 70%. The three-dose compound confirmation screen (1.25, 10, 25 µM) was conducted using the same conditions as in the primary screen.

### The APP reporter-based counter screen

For the APP three dose (1.25, 10, 25 µM) counter dose response assays 293T wild type cells were transiently transfected with three different plasmids coding for the components described. Plasmids for APP and luciferase are described in (Karlström et al., 2002) and the renilla plasmid described above generated for this project was used to normalize firefly values. 24h post transfection cells were seeded on compound plates and cells were cultivated for 24h before luciferase signals were read, as described.

### Dose-response analysis

Compounds for the 11-point (4.00E-05, 2.00E-05, 1.00E-05, 5.00E-06, 2.50E-06, 1.25E-06, 6.25E-07, 3.12E-07, 1.56E-07, 7.81E-08, 3.91E-08M) dose response assays (APP reporter assay and Notch reporter assay) were distributed by two step serial dilutions from compound stocks in master plates using a Bavo automated liquid handling system (Agilent). Two-fold concentrations of compounds were diluted in 10 µl cell culture medium and brought to one-fold by seeding 10 µl containing 4000 cells of the respective cell type on each well of a 384-well plate, as described above. Analogous to the previous assays, all plates contained one DMSO and DAPT control column with the Notch reporter cell line (#A1-11) and a DMSO only and Notch inhibitor control plate was read before the measuring the screening plates. In addition, DAPT and DMSO were spotted in a dose response fashion, and a minimum of four replicates were read. Dose-responses of compounds were recorded in triplicates for each concentration reading out the dual luciferase reporters, as described above. Dose response assays were read together in one session. To measure compound effects, the different assays dose-response curves were fitted using the log(inhibitor) vs. response -- Variable slope (model: Y=Bottom + (Top-Bottom)/(1+10^((LogIC50-X)*HillSlope)) equation. In addition, the area-under-the-curve (AUC) was calculated using Graphpad Prism.

The 14-point dose-response analysis was conducted in a similar fashion, but compounds were spotted on plates via an Echo 650 liquid dispenser instead of diluted from stocks. As described, compounds were sorted into three different baskets corresponding to different dose-ranges (concentration for compounds in basket #1: 4.00E-05, 1.33E-05, 4.44E-06, 1.48E-06, 4.94E-07, 1.65E-07, 5.49E-08, 1.83E-08, 6.10E-09, 2.03E-09, 6.77E-10, 2.26E-10, 7.53E-11, 2.51E-11M, basket #2: 4.00E-05, 2.00E-05, 1.00E-05, 5.00E-06, 2.50E-06, 1.25E-06, 6.25E-07, 3.13E-07, 1.63E-07, 7.50E-08, 3.75E-08, 2.50E-08, 1.25E-08, 4.88E-09M basket #3: 5.00E-06, 1.25E-06, 5.63E-07, 1.88E-07, 6.25E-08, 2.50E-08, 6.88E-09, 2.25E-09, 7.50E-10, 2.50E-10, 8.50E-11, 2.75E-11 1.00E-11, 2.50E-12M). A DMSO plate was read before measuring luciferase activity of compound spotted plates. APP firefly luciferase reporter assays were read with a signal integration time of 0.1 seconds, instead of 0.5 seconds used for the Notch reporter to reduce signal overlap between wells. IC50 values and dose response curves were fitted as described, but the bottom of the assay was defined by the maximum inhibition recorded with DAPT or the GSI LY-411575 (APP counter assay) to define a more precise assay window.

### The C2C12 myogenic assay

8000 C2C12 cells were seeded on gelatin (0.1%, Sigma-Aldrich, cat #G2500) coated cover slides (BD Bioscience) and cultivated in proliferation medium. For gelatin coating, 8-well cover slide chambers (BD Bioscience) were incubated for 60 min with 0.1% gelatin in PBS. The remaining liquid was aspirated, and gelatin was allowed to dry for 24h. Cover slides were stored at 4°C until use. Upon reaching 85% confluence cell culture medium was exchanged to differentiation medium to enhance differentiation into myocytes/myotubes. Cells were then treated with 10 µM of the respective compound for five days and DMSO-treated cells served as a base-line control. Next, cells were washed with 1x cold PBS (ThermoFisher, cat #10010023) fixed with 4% paraformaldehyde (PFA, Sigma-Aldrich, cat #1.00496) in PBS for 10 min at room temperature and washed again with 1x PBS. Fixed cells were either stored at 4°C in PBS or directly processed for immunocytochemistry staining: For immunocytochemistry, cells were permeabilized for 10 min in 0.1% TritonX-100 / PBS, then washed in PBS and unspecific bindings sites were blocked in protein free blocking solution (DAKO, cat #x0909) for 30 min at room temperature, followed by two times washing in PBS. Then, cells were incubated with primary antibody MYH4 (ThermoFisher) in blocking solution and 0.1% TritonX-100 in PBS for 1h at room temperature. Next, cells were washed three times in PBS and incubated with secondary Alexa fluorophore coupled antibody (Invitrogen) in a 1:400 dilution. Labeling of F-actin filaments was conducted by phalloidin-TRITC (Sigma-Aldrich, cat # P1951) in a 1:2500 dilution. Cells were washed three times in PBS and incubated with DAPI (BD Bioscience cat # 564907) for 5 min at room temperature to stain DNA/nuclei. Finally, slides were fixed with mounting medium (ProLong Antifade mounting medium, ThermoFisher, cat #P36934). Bright field and immunocytochemistry images were taken using the SP8 LSM (Leica) at different magnifications and images were recorded as Z-stacks. Images were processed using the FijI (ImageJ) software. As controls, wells with and without differentiation medium were also stained to set the baseline and contrast the differences and stainings of cells with only secondary antibodies were used as negative controls to assess cross-reactivity and background staining. For calculating the MYH4 median fluorescence intensity (MFI) images have been converted into 8-bit greyscale and a standard color threshold was applied (ImageJ). Next, MFI signals per area were measured using ImageJ and divided by the median background intensity (MBI). The nuclei fusion index was calculated as the ratio of nuclei fused per area divided by the total number of nuclei in the field. A minimum of three images has been used for the calculations and two replicate experiments were conducted.

### Three dose response gene expression analysis

Adherent breast cancer cells were seeded in 24-well cell culture plates (ThermoFisher, Nunc, Delta surface, cat #144530) and T-ALL cells growing in suspension (Jurkat, RPMI-8402) were seeded in non-adherent 24-well plates (ThermoFisher, Nunc, cat #144530). 60.000 cells were seeded and treated with 0.1, 1 and 10 µM of each compound. As controls, CB-103 (a non-GSI Notch inhibitor) and the GSI RO4929097 were used with the same dose regimen. Cells were harvested 24h post treatment and RNA isolation and gene expression analysis (qRT-PCR) was performed as described above.

### Growth inhibition in tumour cell lines

Metabolic activity in different tumour cell lines was assessed in 14-point dose response regimens (5.00E-05, 2.50E-05, 1.25E-05, 5,00E-06, 2.50E-06, 1.25E-06, 5.00E-07, 2.50E-07, 1.25E-07, 5.00E-08, 2.50E-08, 1.25E-08, 5.00E-09, 2.50E-09 M). Compounds were spotted on white 384-well tissue culture plates (Greiner, cat #781080) using an Echo 525 Acoustic liquid handler (Labcyte). Cells were grown to 80% confluence, detached if required, counted and 20 µl containing 2.000-4.000 cells per well was then seeded in each well and cultivated for three (4.000 cells seeded) and six days (2.000 cells seeded). As a control, a minimum of 24 wells containing cells incubated with only DMSO equimolar to the highest compound concentration or the GSI RO4929097 or the Notch inhibitor CB-103 were included. Luciferase activity (reading out metabolic activity) was determined using the CellTiter-Glo 2.0 kit (CTG, Promega, cat #G9242) following the manufacturer’s instructions and luminescence was measured on a SpectraMaxi3 plate reader (Molecular Devices).

### Subcellular fractionation experiments

MDA-MB-231 cells were grown to 80% confluence in T-75 cell culture flasks (Sarstedt, cat # 83.3911.002) and compounds were directed diluted in cell culture medium to a concentration of 2 µM. Cells were then incubated for 24h and cell fractionation experiments were performed using a cell fractionation kit (Cell signalling, cat #9038S), following the manufacturer’s instructions.

### Immunocytochemistry

20.000 MDA-MB-231 cells were seeded on cover-slides (BD Bioscience) and allowed to attach, and the next day they were treated with 5 µM of compounds for 24h. Subsequently, immunocytochemistry staining to localize NOTCH1 was carried out as described above for C2C12 cells. Cells were incubated with primary EEA1 antibody at 1:200 dilution (BD Bioscience, cat #610457) and a NOTCH1 antibody was used at 1:100 dilution (Cell signalling, cat #D6F11). Images were taken using the SP8 LSM (Leica) with a 40X objective and were further processed using Fiji (ImageJ) software. A minimum of four images per treatment was recorded.

### Cycloheximide chase experiments

293T Notch reporter cells (clone #A1-11) were grown to 80% confluence, detached and counted. 8.000 cells were then seeded in each well of a 96-well tissue culture plate (Sarsted, cat # 83.3924). The next day, 10 µM of the respective compound and 100 µg/ml Cycloheximide (CHX) solution (Sigma-Aldrich, cat #C4859-1ML) were added. Each well was then incubated with 50 µl of compound and CHX solution and compound and CHX solution were added after the indicated intervals to another well for a total time of eight hours. Next, medium was aspirated off and cells were lysed in RIPA buffer (Thermofisher, cat #89900) containing Protease and Phosphatase inhibitors (Thermofisher, cat #78442) for 30 min on ice. Lysates were thoroughly vortexed, centrifugated for 10 min at >13.000x g and supernatant was transferred to a fresh cup and stored at -80°C.

### DHODH inhibitor and uridine rescue experiment

293T Notch reporter cells (#A1-11) were grown to 80% confluency, detached and mixed with the indicated compounds and doses with or without uridine (Sigma-Aldrich, cat #U3003). 10.000 cells/well were then seeded into 96-well cell culture plates (ThermoFisher, cat #136102) and incubated for 24h. DMSO treatment with or without uridine served as control for each treatment. Cells were then subjected to Notch reporter assays, and luciferase signals were recorded on a Varioscan Lux microplate reader (ThermoFisher). Reporter signals from each treatment were normalised to the respective DMSO control, as described above and three independent assays were recorded with at least two technical replicates per treatment.

### Surface Biotinylation experiment

For labelling of membrane proteins, MDA-MB-231 cells were grown to 80% confluency in 6-well cell culture dishes (ThermoFisher, cat #104675) and treated with 2 µM of the indicated compounds for 24h. Plates were then put on ice for 10 min to slow down cellular metabolism and halt internalisation of membrane proteins. Next, 0.5 mg/ml Sulfo-NHS-Biotin (ThermoFisher, cat #21217) in PBS was added to every second well for 40 min and non-treated wells served as controls. Excess of biotin was removed by washing the plates with PBS followed by incubation with 100 mM glycin (Sigma-Aldrich, cat #G8898) in PBS on ice. Wells were washed twice with ice-cold PBS and next, 500 µl lysis buffer was added to each well (50 mM Tris-HCl pH 7.4, 150 mM NaCl, 1% Triton X-100, (Sigma-Aldrich, cat #X100), 2 mM EDTA, including protease and phosphatase inhibitors, Thermofisher, cat #78442). Lysates were thoroughly vortexed, centrifugated at 14.000xg for 10 min at 4°C and 50 µl of each supernatant was taken as input fraction and stored until further use at -80°C. NeutrAvidin Agarose beads were pre-equilibrated in lysis buffer for 5 min and 50 µl was added to each sample. Pull-down fractions were incubated on an end-to-end rotator over night at 4°C. Next, beads were washed three times in lysis buffer for at least 10 minutes per wash and dry beads were mixed with 4x Laemmli buffer (Bio-rad, cat #1610747) and stored at -80°C until use.

### HiBit reporter assay

20 µl containing 4.000 cells were seeded on 384-well pre-spotted compound plates (Greiner, cat #781080). Cells were incubated for 24h and viability was determined using CellTiter-Fluor (Promega, cat # G6080), following the manufacturer’s instructions. Next, cells were subjected to the HiBit assay according to the manufacturer’s instructions (Promega, cat # N3030). Prior to cell lysis, the tissue culture plates were allowed to cool down to room temperate. Fluorescence and luminescence signals were recorded using a SpectraMax i3 (Molecular devices) and signals were read at different time-points to determine optimal signal stability. Cells cultivated only with DMSO were used as a reference to determine effects of compounds. MG132, Bafilomycin A1 and the GSI RO4929097 served as additional controls. Celastrol was used to determine the assay window, and a minimum of two technical replicates were measured per assay.

### Western blot analysis

Cells were washed in ice-cold PBS and incubated for 30 min on ice with RIPA cell lysis buffer (Thermofisher, cat #89900) containing protease and phosphatase inhibitors (Thermofisher, cat #78442), scraped off and homogenized using a 25G syringe, and subsequently centrifugated for 10 min at >13.000xg. The supernatant was mixed with 4x Laemmli buffer (Bio-rad, cat #1610747) and heated for 5 min to 95°C and either loaded on MiniProtean Tris-Glycine TGX gradient gels (Biorad) or stored at -80°C until further use. Protein lysates were separated on MiniProtean 4-15% or 4-20% Tris-Glycine TGX gradient gels (Bio-rad). After electrophoresis, gels were placed directly onto a TransBlot Turbo PVDF or nitrocellulose membrane (Bio-rad, cat #1704157 and cat #1704159) and transferred using the “HIGH MW” protocol (Bio-Rad). Subsequently, membranes were blocked for 1h at room temperature using Clear Milk Blocking Buffer (ThermoFisher, cat #37587) supplemented with 0.1% Tween20 (Sigma-Aldrich, cat #P1379). Detection of antibody binding was performed by using Clarity Western ECL-Substrate (Bio-rad, cat #1705060S). Beta-Actin protein signals were used as a loading control. Western blot membranes were in some experiments stripped after development (indicated in the figure in each respective case) by washing the membranes twice for 10 min in PBS, followed by incubation in membranes in western blot stripping buffer (ThermoFisher, cat #21059) for 30 min at room temperature and gentle agitation. Stripped membranes were washed twice with PBS and incubated for 30min in blocking buffer before incubation with primary antibody. A list of all antibodies used in this study is provided in Supplemental Table 6.

## Acknowledgements

We kindly thank Sven Poetsch, Lisa Alberts and Eva Hambruch at Merck Healthcare KGaA, Darmstadt, Germany for assistance and support at various stages of the screening project. We also thank the Chemical Biology Consortium Sweden (CBCS) team at SciLifeLab, Stockholm, Sweden (Hannah Axelsson, Simon Moussad and Francesco Massai), for conducting the large-scale screening. Further, we thank Francesca del Gaudio and Ellinor Langels for support with the C2C12 and uridine assays, respectively. We are also grateful for the technical support from the single-cell core facility (SICOF), Huddinge, Sweden, for spotting of the individual compounds.

## Author contributions

UL, EBB, DW and AS conceived the study and designed the experiments. EBB and MH performed the experiments. Data were collected and analysed by EBB, DW, AS, TH, SL and UL. The manuscript was written by UL and EBB, with input from DW, AS, MH and SL, and reviewed by all authors. All authors have read and approved the final version of the manuscript.

## Supplemental Figures

**Supplemental Figure 1:**

**Establishing the Notch reporter assay in the 384-well format. A)** The Notch reporter (firefly) and control (renilla) luciferase activities were analysed at different timepoints (10 and 25 min for firefly and 1, 10 and 40 minutes for renilla), as indicated. The effect of DMSO versus DAPT was also analysed, as indicated. **(B)** The Notch reporter (firefly) and control (renilla) luciferase activities were analysed at different concentrations of DMSO (0.0125 to 2%) and DAPT (3 to 500 mM). The dotted line indicates the selected concentration of compounds and controls for the primary screen, 10 µM. Firefly and renilla signals are presented at the top and at the bottom the firefly/renilla luciferase ratio (left) and Z-factor of firefly, renilla and firefly/renilla ratio (right) is presented. **(C)** Signal intensities for firefly and renilla luciferase signals across DMSO plate at different timepoints, as indicated. Row numbers from A-P, column numbers from 1-23. At the bottom the firefly/renilla luciferase ratio is presented.

**Supplemental Figure 2:**

**Validation of the Notch reporter assay. (A)** Analysis of the effects of FLI-06, LY-450139 (Semagacestat), LY-411573 and DAPT at three doses (1, 5, 10 mM) in the Notch reporter assay. **(B)** The signal-to-background ratio of DMSO and DAPT controls across 120 plates is shown for firefly and for renilla luciferase individually and for the normalised ratio of DMSO and DAPT. **(C)** Analysis of control signals for firefly and renilla luciferase in the presence of DMSO or DAPT across 120 plates. The signals after normalization are shown below to the right. **(D)** Average (Fluc/Rluc) ratio across 120 plates. To the right, the DMSO/DAPT ratio in percent of DMSO controls across 120 plates is presented. **(E)** The Z’-factor for firefly luciferase activity and the Z’-factor for the ratio between DMSO and DAPT is shown. **(F)** The Coefficient of variation across 120 plates is shown for firefly and for renilla luciferase individually and for the normalised ratio of DMSO and DAPT. **(G)** Analysis of three luciferase inhibitors (PTC-124, luciferase inhibitor 1 and Nano-Luc inhibitor) as well as FLI-06, LY 450139, LY 411573 and DAPT in the Notch reporter assay at three different dosages (1.25, 10 and 25 mM). **(H)** The signal-to-background ratio of DMSO and DAPT control is shown for firefly and for renilla luciferase individually and for the normalised ratio of DMSO and DAPT. **(I)** The Z’-factor for firefly luciferase activity and for the ratio is shown. **(J)** The coefficient of variation is shown for firefly and for renilla luciferase and for the normalised ratio of DMSO and DAPT. **(J)** Left, average firefly and renilla luciferase signals of DMSO and DAPT controls is shown and averaged DMSO and DAPT ratio is shown to the right. **(C)** Reporter signals of DMSO and DAPT after normalization is shown to the left and the DMSO/DAPT ratio relative to the DMSO controls is shown to the right. The analysis was carried out using a total of six compound plates.

**Supplemental Figure 3:**

**Validation of the APP-C99 counter assay. (A)** Signal intensities for firefly (upper) and renilla (lower) luciferase signals across DMSO plate. Row numbers from A-P, column numbers from 1-22. At the bottom the firefly/renilla luciferase ratio is presented. The red dots indicate the signals of the Notch-reporter, that was used for comparison. **(B)** Analysis of three luciferase inhibitors (PTC-124, luciferase inhibitor 1 and Nano-Luc inhibitor) as well as FLI-06, LY 450139, LY 411573 and DAPT in the APP-C99 counter assay at three different dosages (1.25; 10 and 25 mM). **(C)** Analysis of data from the APP-C99 counter assay and the Notch confirmation assay to remove potential GSIs decreasing reporter signal to less than 70% in the counter assay at three different dosages of compound (1.25, 10 and 25 µM). The blue box marks compounds that met the criteria and were further investigated, while the red box presents compounds that failed to meet the thresholds and were sorted out. **(D)** Analysis of LY 450139, LY 411573 and DAPT in the APP-C99 screen at four different dosages (3.125; 6.25; 12.5 and 25 mM), and of luciferase inhibitor 1 and PTC-124 at 50 mM. **(E)** Dose-response curves from the Notch reporter assay (red) and the APP-C99 assay (blue) using compounds at 11 different doses. **(F)** Plate statistics for the 11 does-points APP-C99 and Notch reporter screens are shown, as indicated. **(G)** Left, average firefly luciferase signals of DMSO and DAPT controls are shown and averaged DMSO and DAPT ratio is shown to the right and below the respective averaged renilla luciferase signals are shown. The reporter signals of DMSO and DAPT after normalization are shown at the bottom, left and the DMSO/DAPT ratio relative to the DMSO controls is shown at the bottom, right.

**Supplemental Figure 4:**

**Validation of the APP-C99 counter assay in a 14-point dose-response regimen. (A)** Dose-response curves from the APP-C99 counter assay in a pilot 9-point and 14-point dose regimen for RO4929097, YO-01027, LY-411575, BMS-983970, DAPT and CB-103, as indicated. Black cell 384-well plates were also evaluated (data not shown). **(B)** Plate statistics for the APP-C99 counter assay in the 14-point dose regimen format. Average control signals, reporter signals after normalization, average Fluc/Rluc ratio in response to DAPT or DMSO, ratio of DMSO/DAPT and Z’-factor were established for the various compounds, as indicated. **(C)** Dose-response curves from the APP-C99 counter assay using compounds at 14 different concentrations. **(D)** Dose-response curves from the Notch reporter assay in a pilot 9- and 14-point dose regimen for RO4929097, YO-01027, LY-411575, BMS-983970, DAPT and CB-103, as indicated. **(E)** Plate statistics for the Notch reporter assay in the 14-point dose regimen format. Average control signals, reporter signals after normalization, average Fluc/Rluc ratio in response to DAPT or DMSO, ration DMSO/DAPT and Z’-factor were established for the various compounds, as indicated. **(F)** Dose-response curves from the Notch reporter assay using compounds at 14 different concnetrations.

**Supplemental Figure 5:**

**Analysis of compounds B and C and respective analogues for growth-inhibitory effects on T-ALL and breast cancer cell lines.** Graphs showing 14-point growth-response curves for the level of metabolic activity in three T-ALL cell lines (JURKAT and TALL) at three and six days after treatment with compound B, C and respective analogues, and CB-103 and RO4929097 as controls, as indicated. **(B)** Graphs showing 14-point growth-response curves for the level of metabolic activity in three breast cancer cell lines (MDA-MB-231, MDA-MB-468 and HCC70) at three and **(C)** six days after treatment with compound B, C and respective analogues and as controls CB-103 and RO4929097, as indicated.

**Supplemental Figure 6:**

**Analysis of Notch downstream gene expression in response to compounds A-E. (A)** Analysis of mRNA expression levels of Notch downstream genes analysed by qPCR, as indicated. Values in response to three different doses (0.1; 1 and 10 mM) of compound A-E, and as controls CB-103 and RO4929097, in JURKAT (upper) and RPMI-8402 (lower) cell lines are shown. **(B)** Corresponding analysis as in **(A)** for MDA-MB-231 (upper), MDA-MB-468 (lower) cell lines are shown. Plots with error bars represent mean ± SD of two biological replicates. Bars without error bars indicate that only technical replicates (n=1) were available.

**Supplemental Figure 7:**

**Establishing conditions and validation of the HiBiT assay. (A)** Luciferase activity in the HiBiT assay for two different HiBiT clones (#E8 and #F2) and HiBiT control clone #D2 (empty vector) 20 minutes (upper) or 40 minutes (lower) after lysis. The Z-factors for control cell line #D2 and HiBit clones #E8 and #F2 are also presented **(B)** Analysis of luciferase activity from subclone #A4 in response to compounds A-E, CB-103, RO4929097 and MG132 for 24h at 10 mM concentration. The HiBiT control clone #D2 was included as control. Outliers were removed using ROUT (Q=1%), as indicated. **(C)** Graphs showing 14-point dose-response curves for the level of HiBiT luciferase activity in the HiBiT reporter cell line of two averaged experiments 24h after treatment with compound B and C at dose-ranges from 50 mM to 25 nM. In addition, dose-response curves for cell viability (labelled CTF Viab; using the CellTiter-Fluor Viability assay, CTF) are shown. RO4929097 and CB-103 were used as Notch inhibitor controls, Bafilomycin and MG132 were used to block protein degradation and Celastrol was as a control to induce apoptosis. Data shown for compounds B and C is also depicted in Figure 7H.

## Supplemental Tables

**Supplemental Table 1:**

List of 189 potential inhibitor compounds and levels of inhibition in the Notch reporter assay at three different doses: 1.25; 10 and 25 µM. Ranking of inhibitors compared to the primary screen.

**Supplemental Table 2:**

List of potential inhibitor hits and their response in the APP-C99 counter assay in comparison to the 3-point dose-confirmation assay at three different doses: 1.25; 10 and 25 µM.

**Supplemental Table 3:**

List of 39 potential inhibitors after screening through an 11-point dose-response APP-C99 counter-assay

**Supplemental Table 4:**

List of structures for 16 candidate inhibitor hits and 47 analogues screened in the APP-C99 counter-assay reporter screen using a 14-point dose-response regimen.

**Supplemental Table 5:**

a) List of 52 (15 candidate inhibitor hits and 37 analogues) and their response to the Notch reporter screen using a 14-point dose-response regimen.

b) List of potential inhibitor hits and their response in CellTiterGLO and HiBit assays.

**Supplemental Table 6:**

List of all antibodies and primers used in this study.

