## Supplemental Figures 1-7 for "Disrupting Notch signalling by a small molecule inhibiting dihydroorotate dehydrogenase activity"

### Supplemental Figure 1

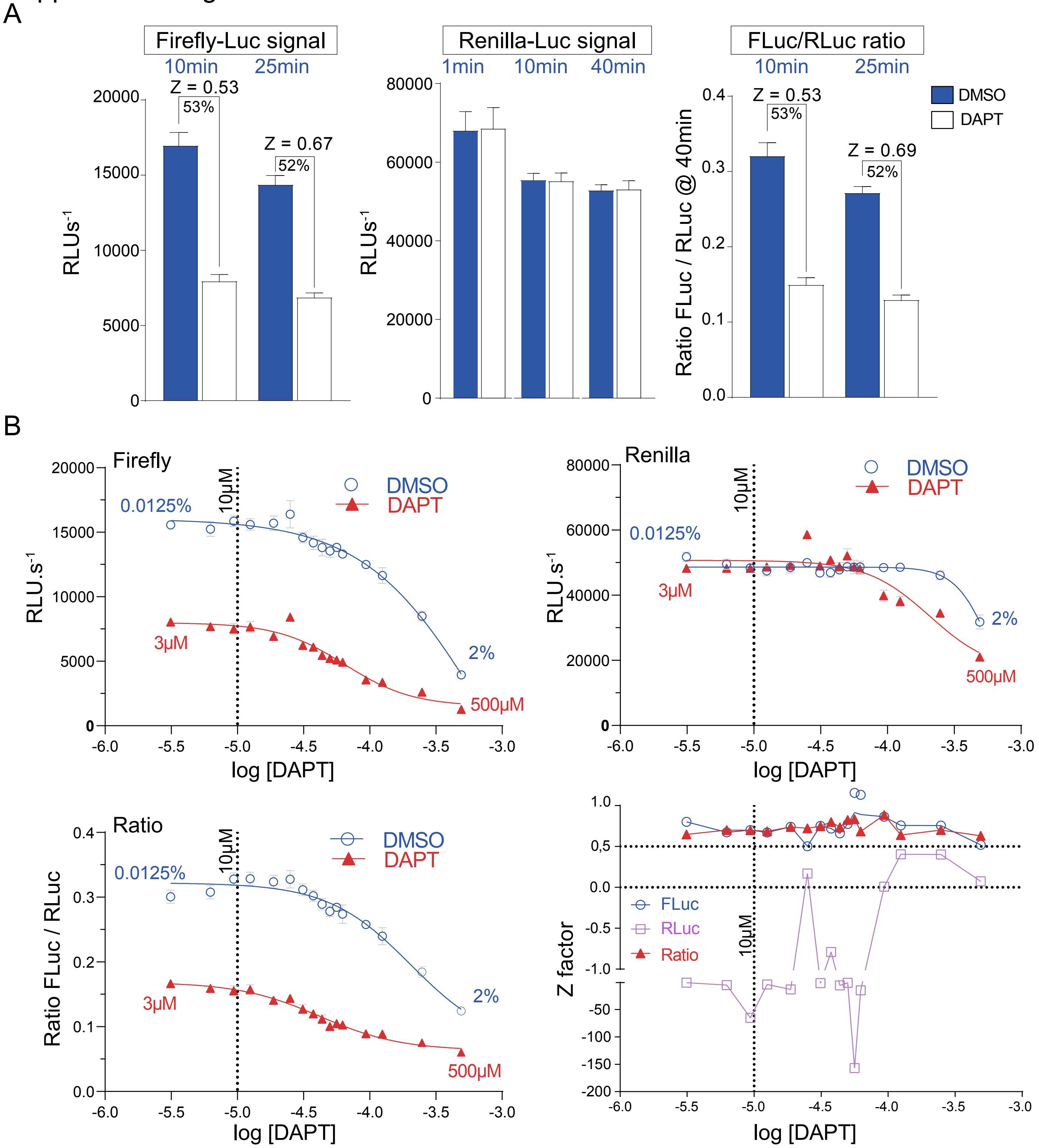

### Supplemental Figure 1 continued
C

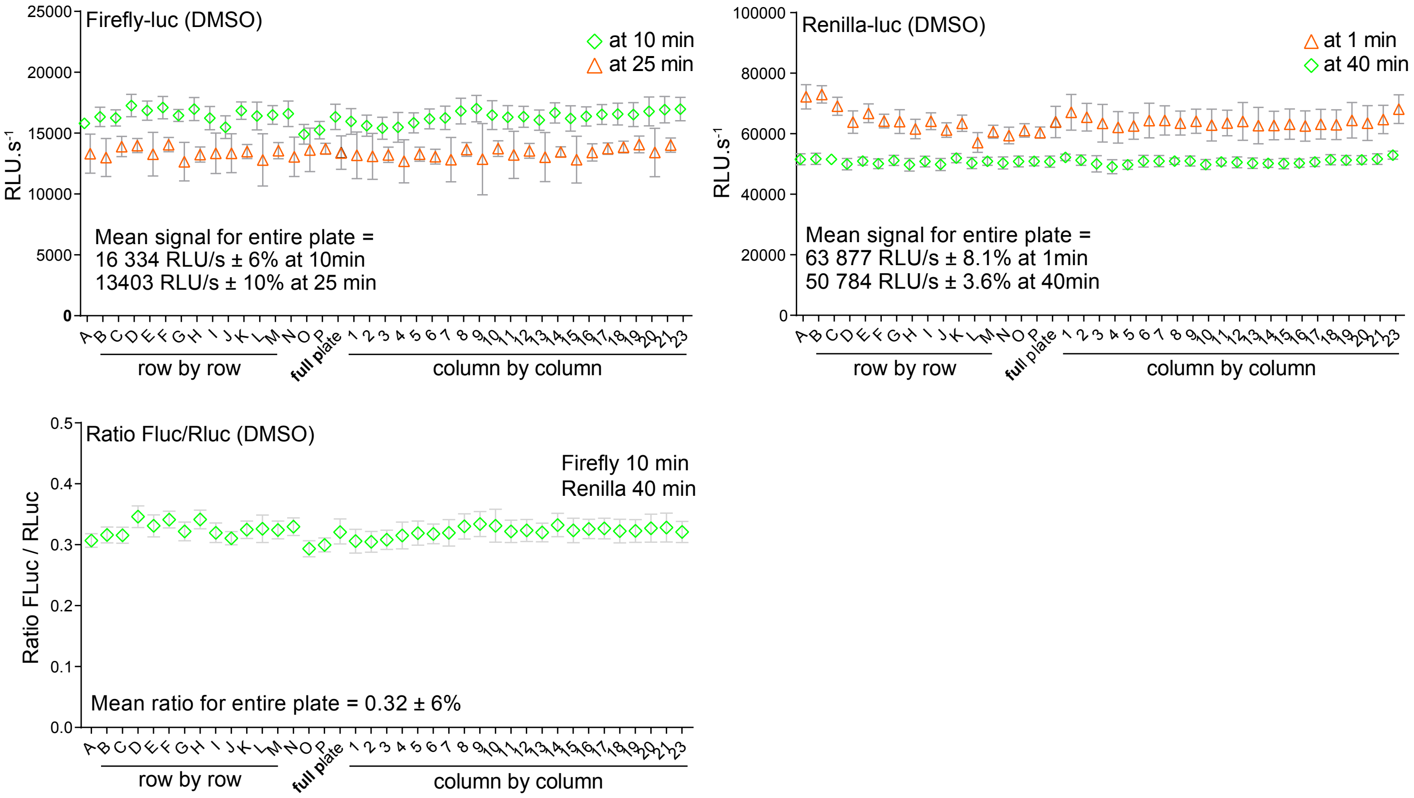

### Supplemental Figure 2

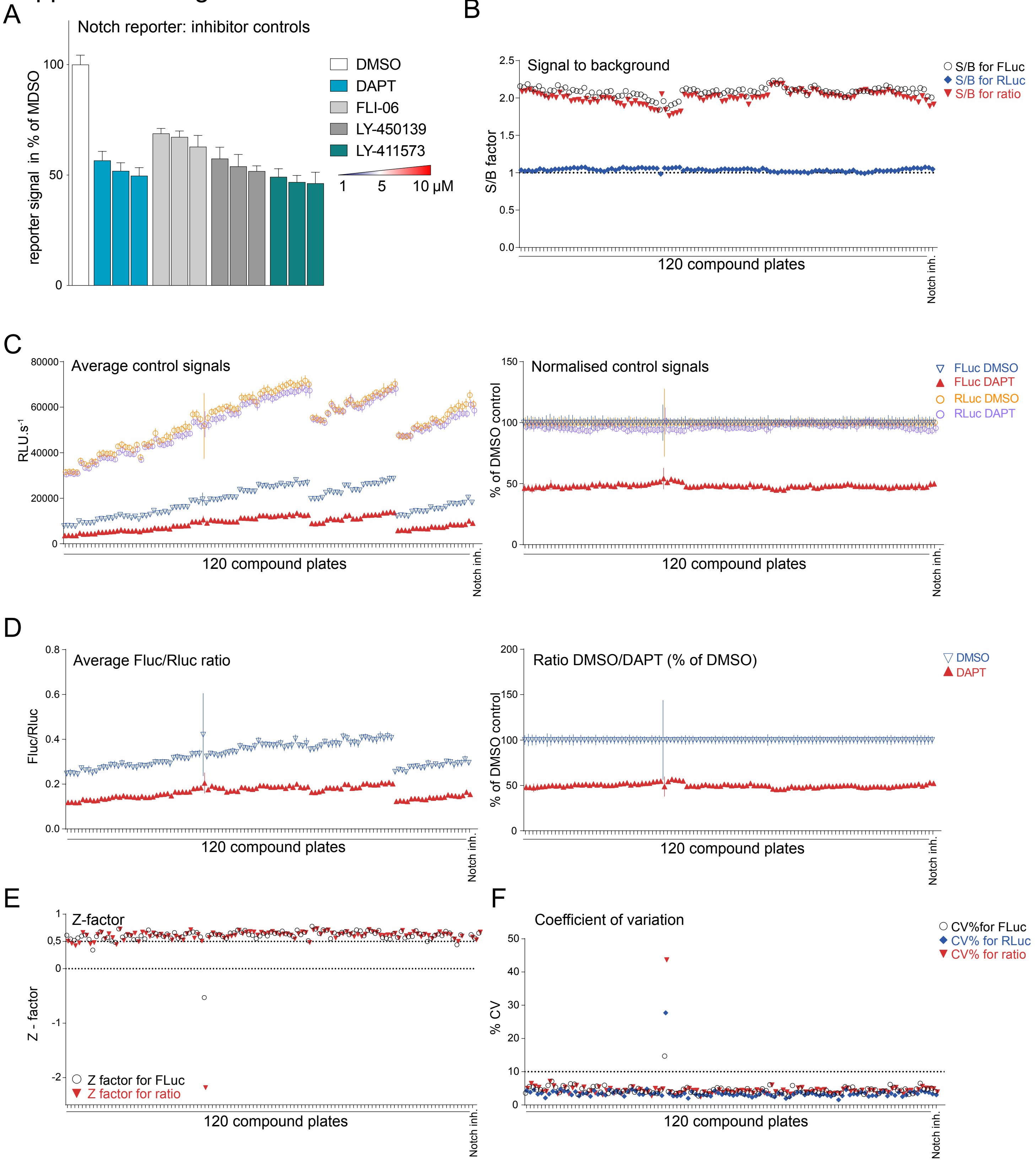

### Supplemental Figure 2 continued

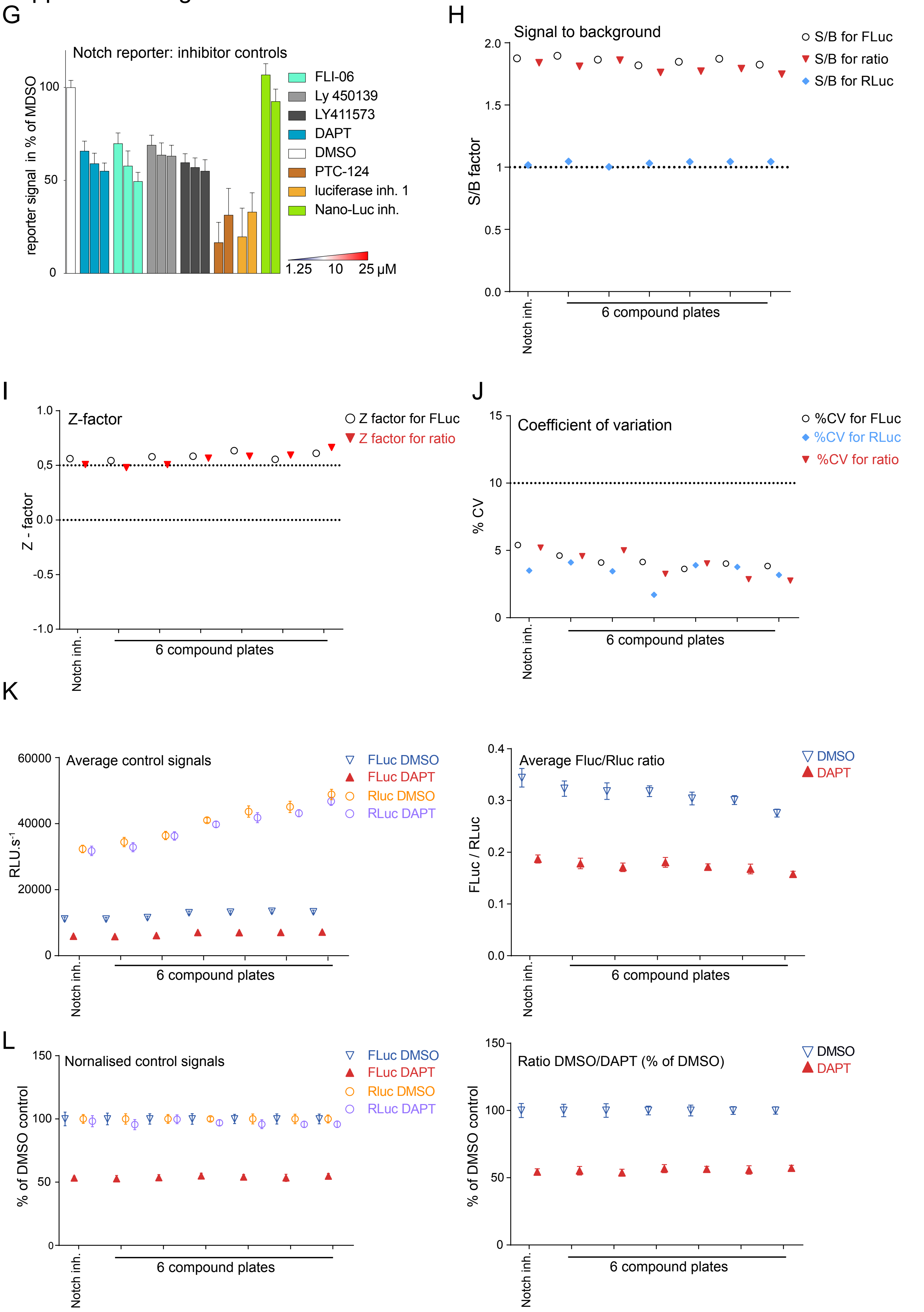

### Supplemental Figure 3

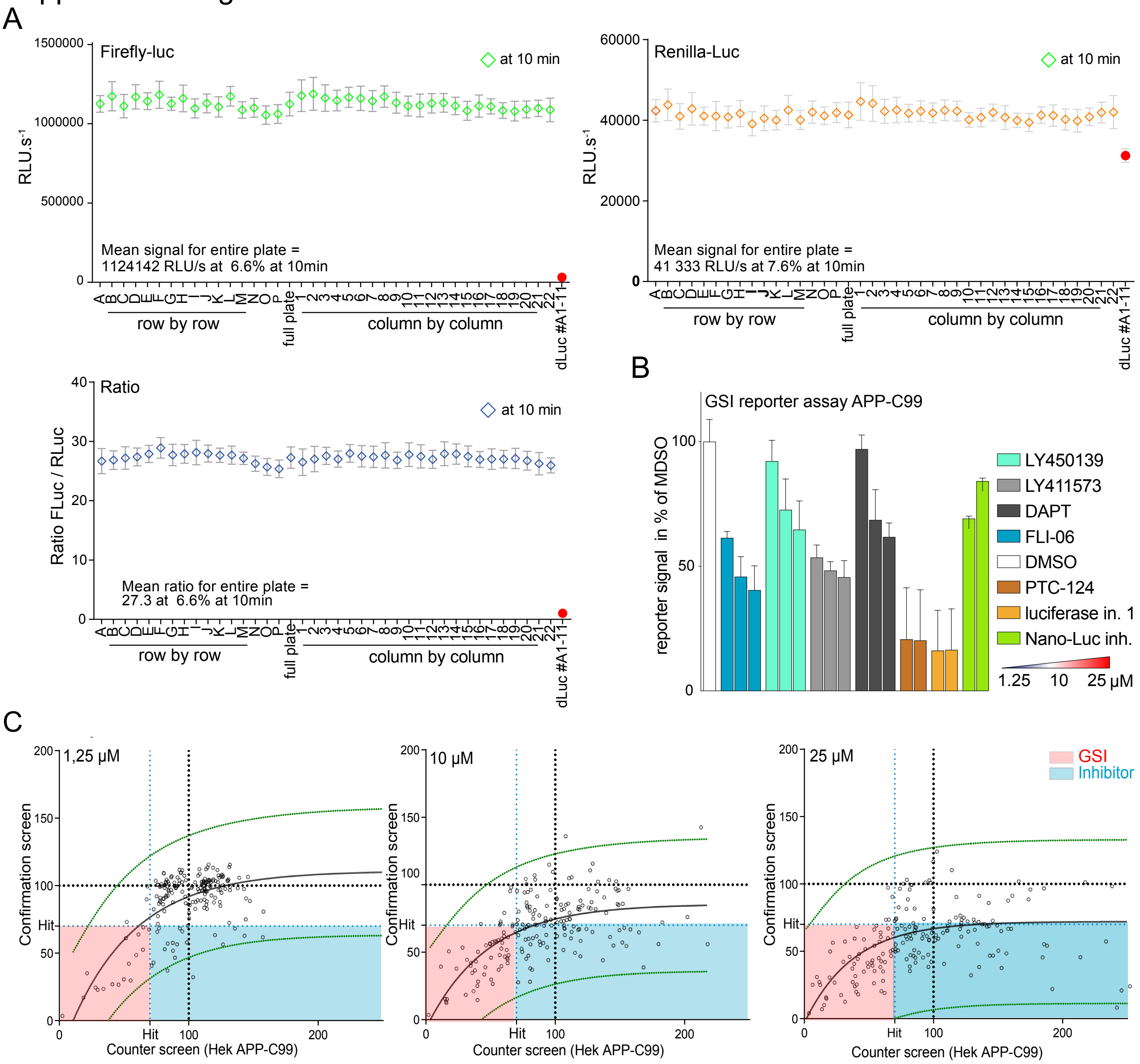

### Supplemental Figure 3 continued

D

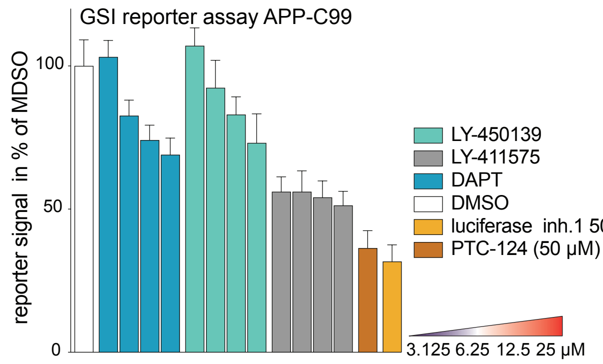


E

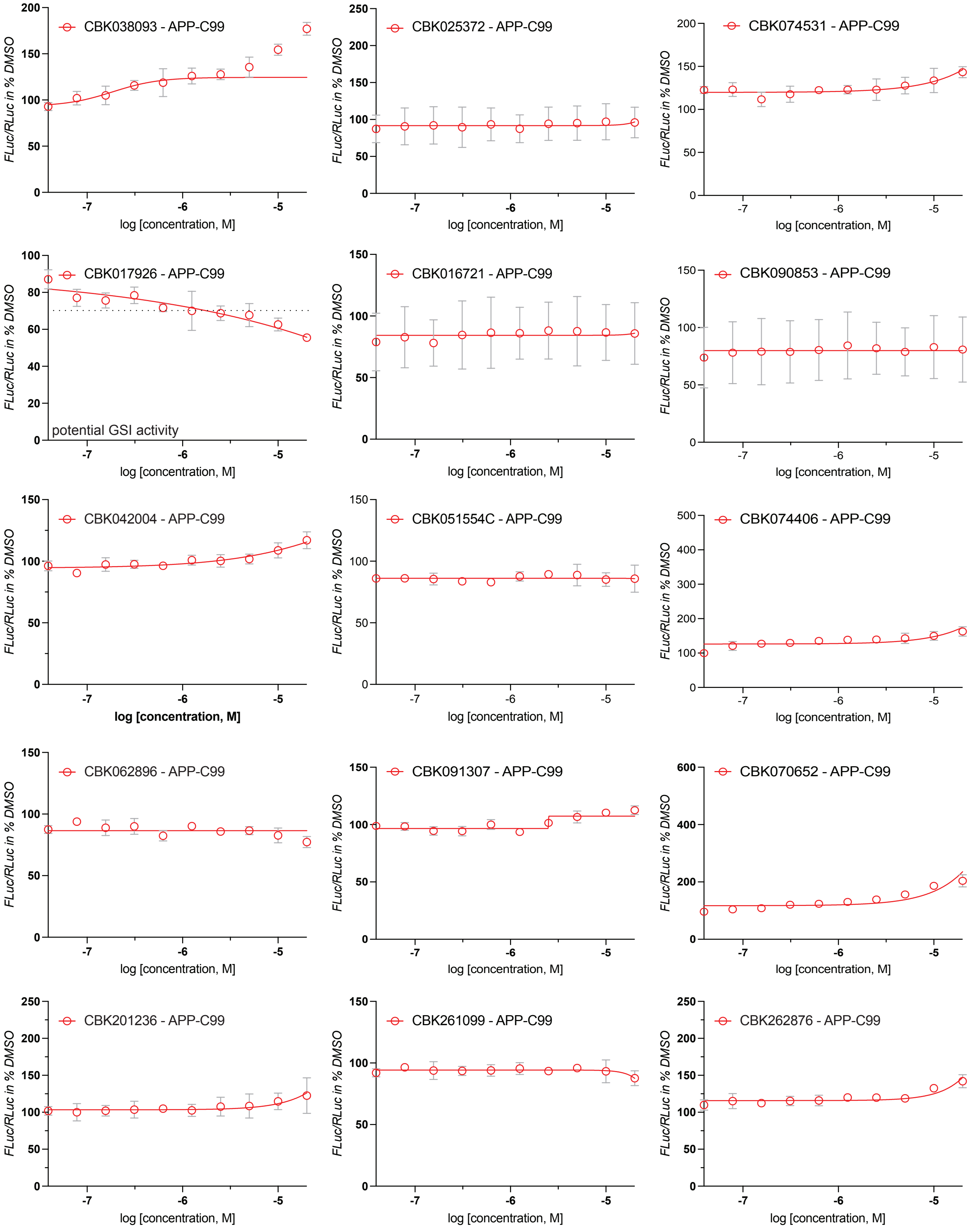

### Supplemental Figure 3E continued

#### 11-point dose-response curves: APP-C99 counter-assay

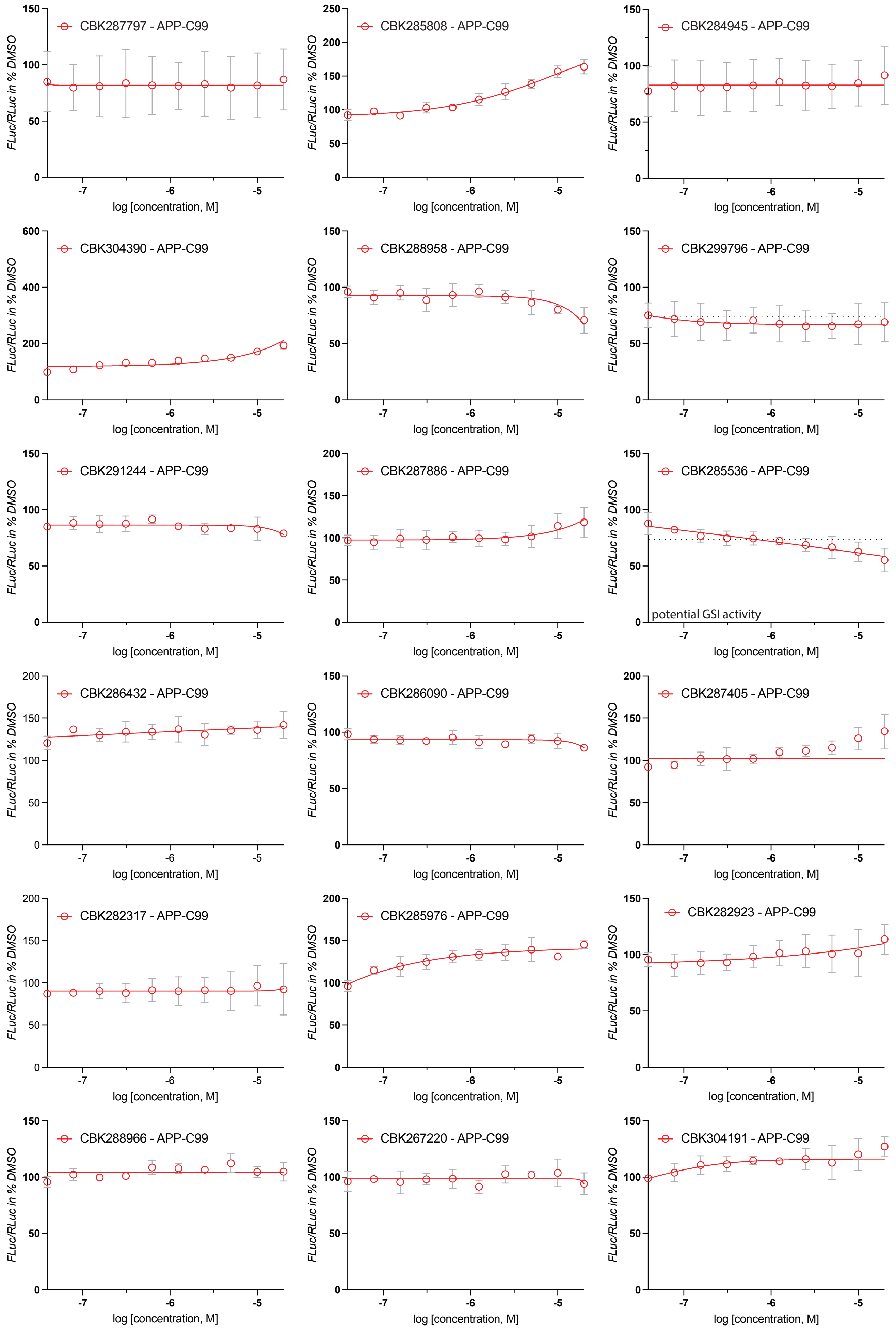

### Supplemental Figure 3E continued

11-point dose-response curves: APP-C99 counter-assay

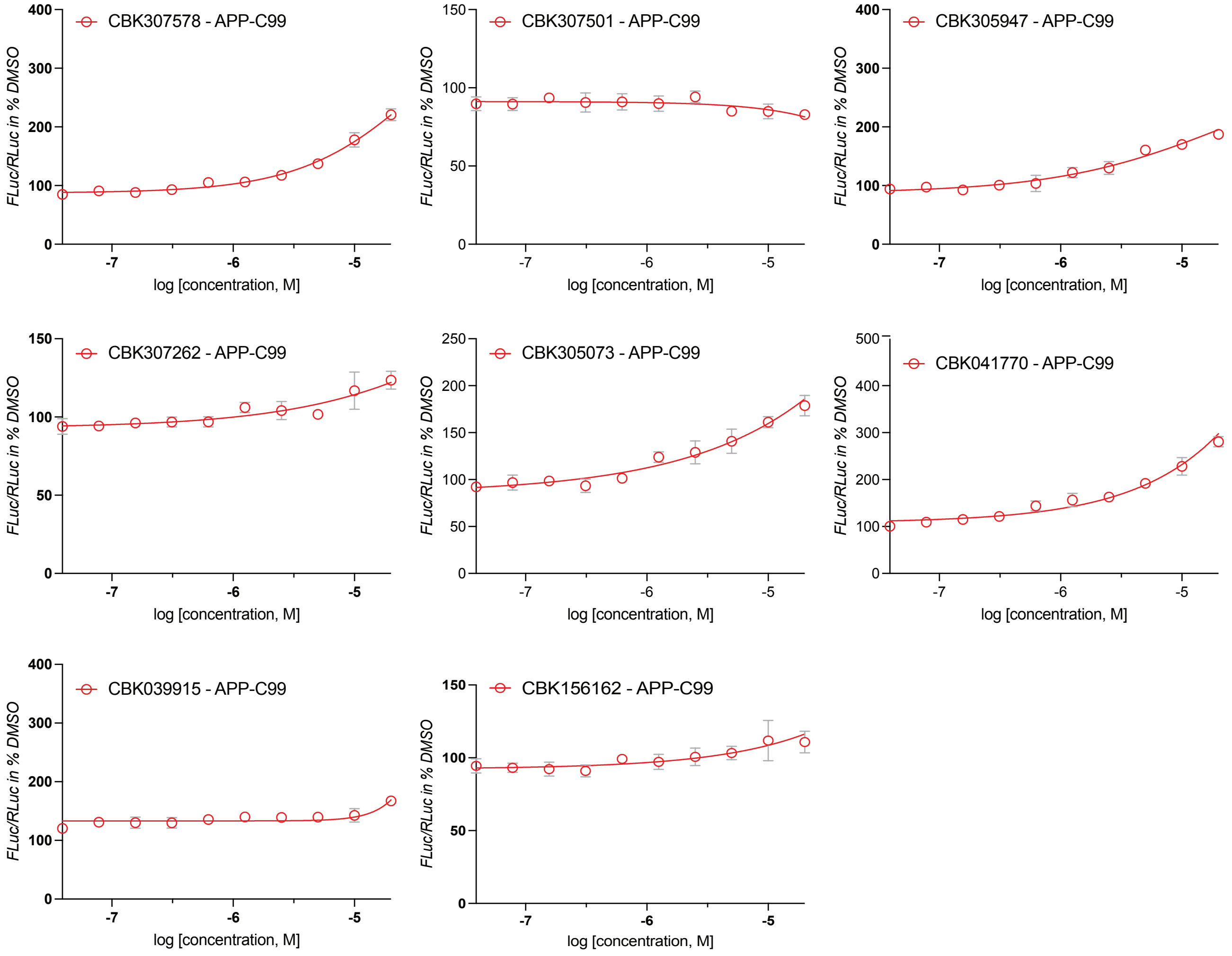

### Supplemental Figure 3 continued

F

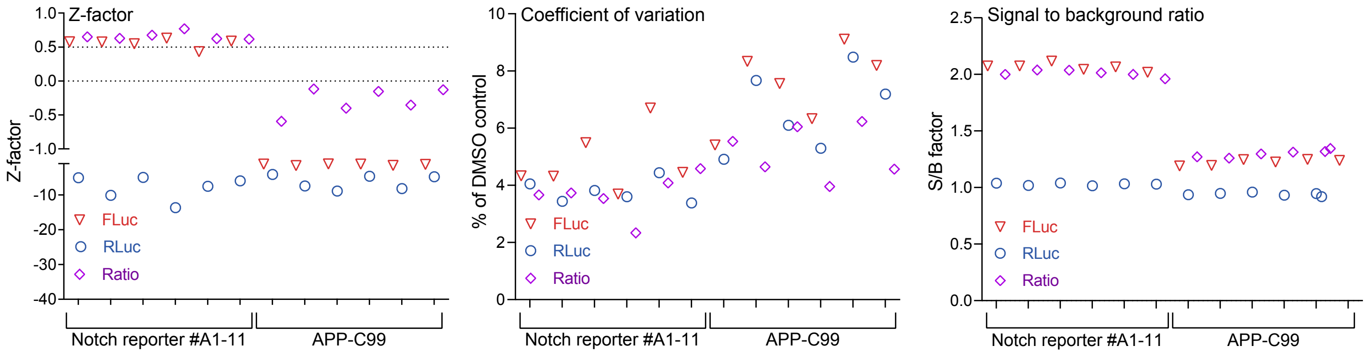


G

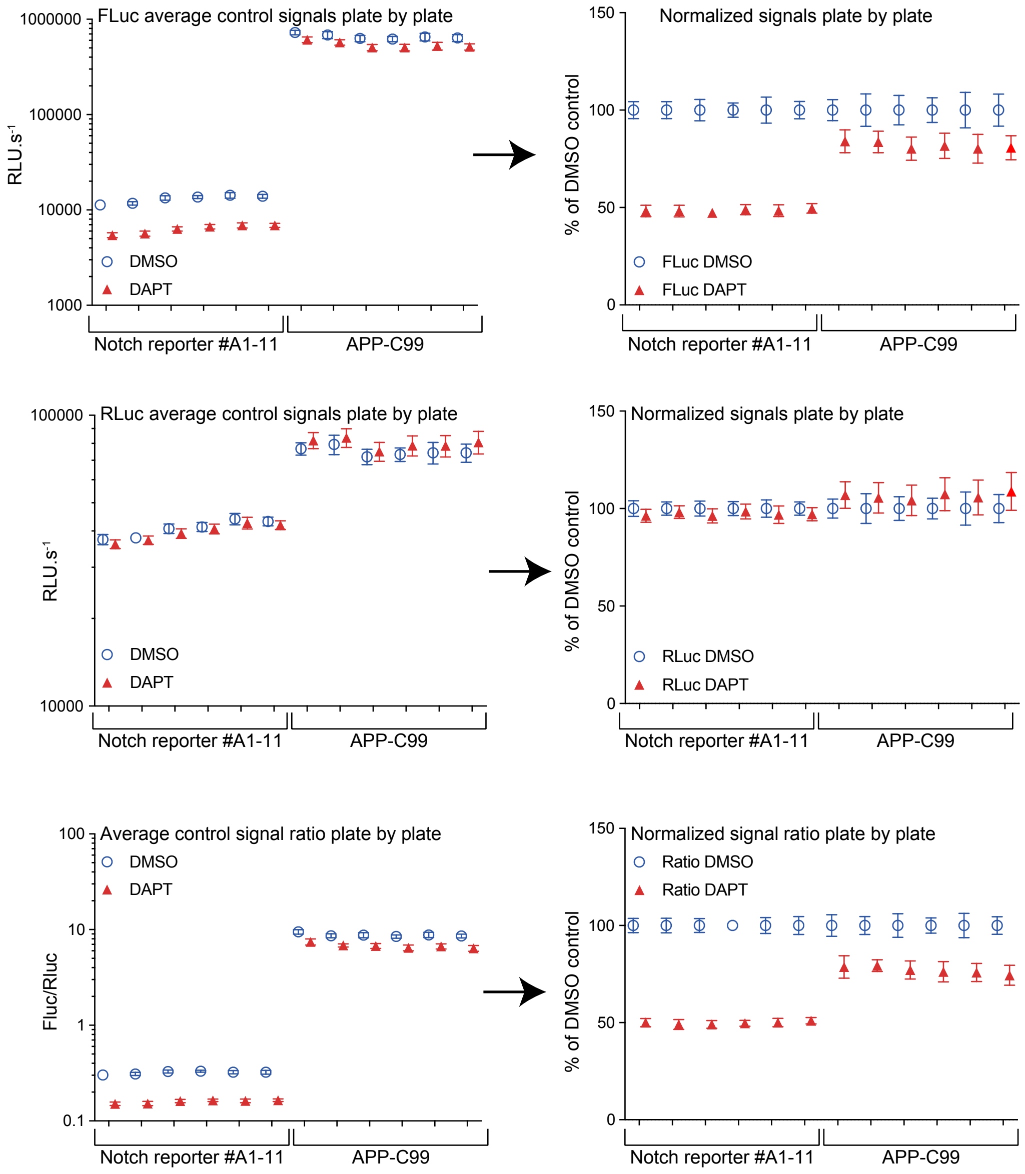

Supplemental Figure 4

A

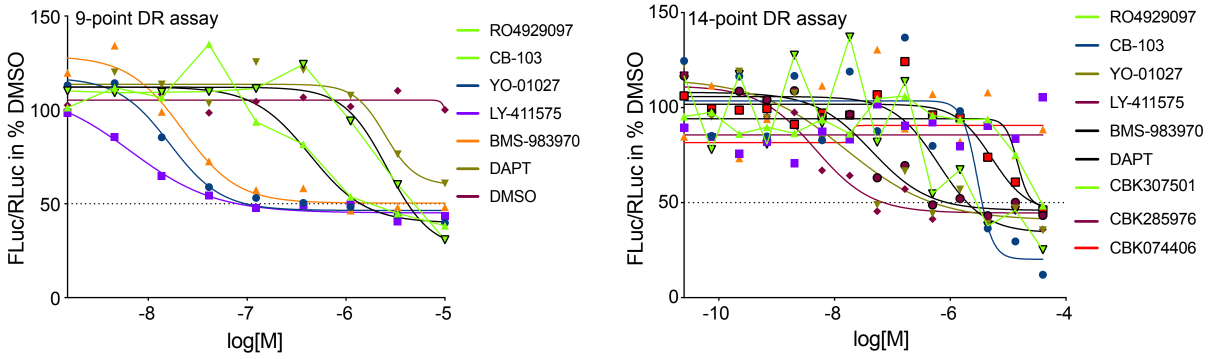


B

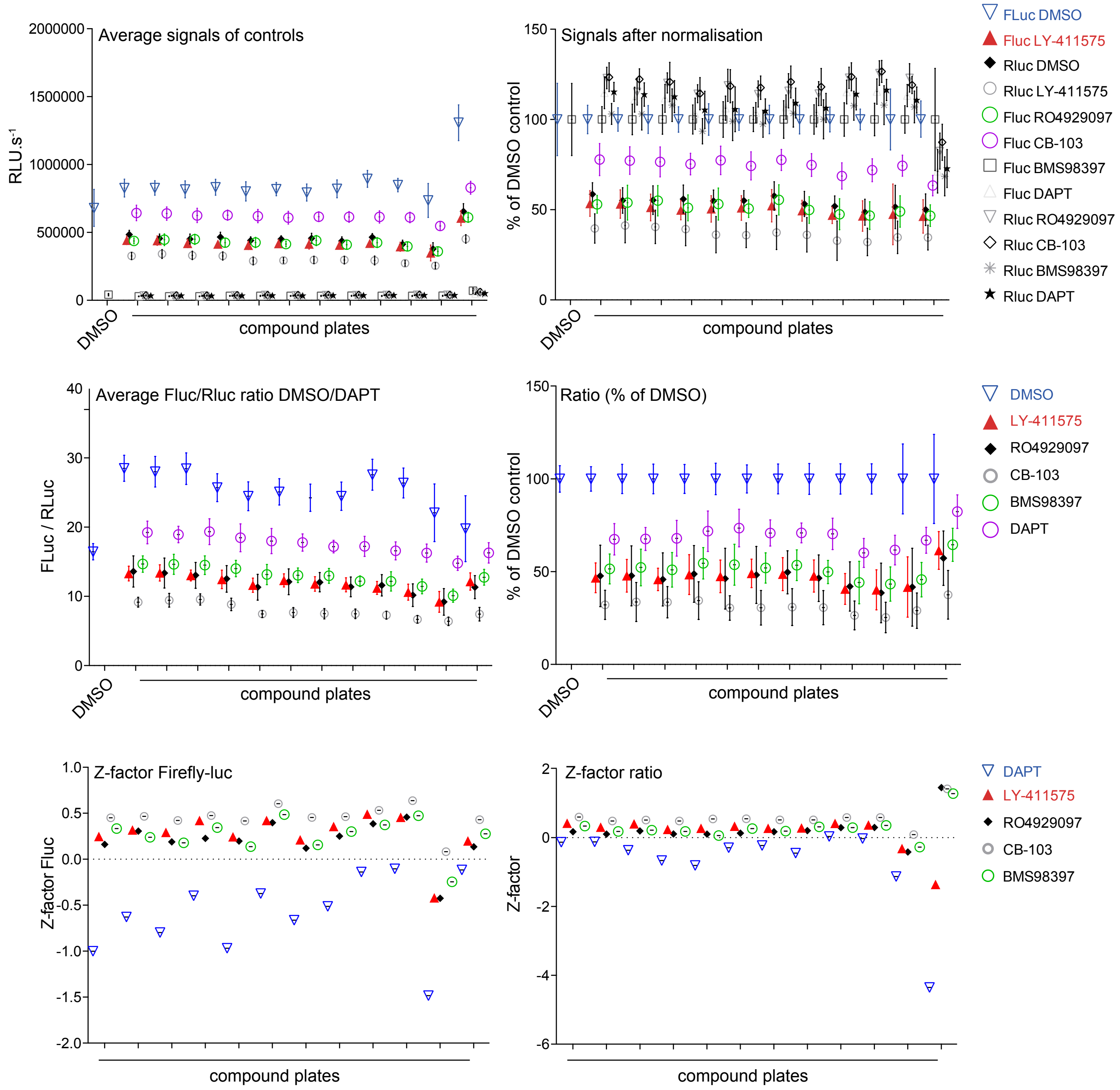

Supplemental Figure 4 continued

C

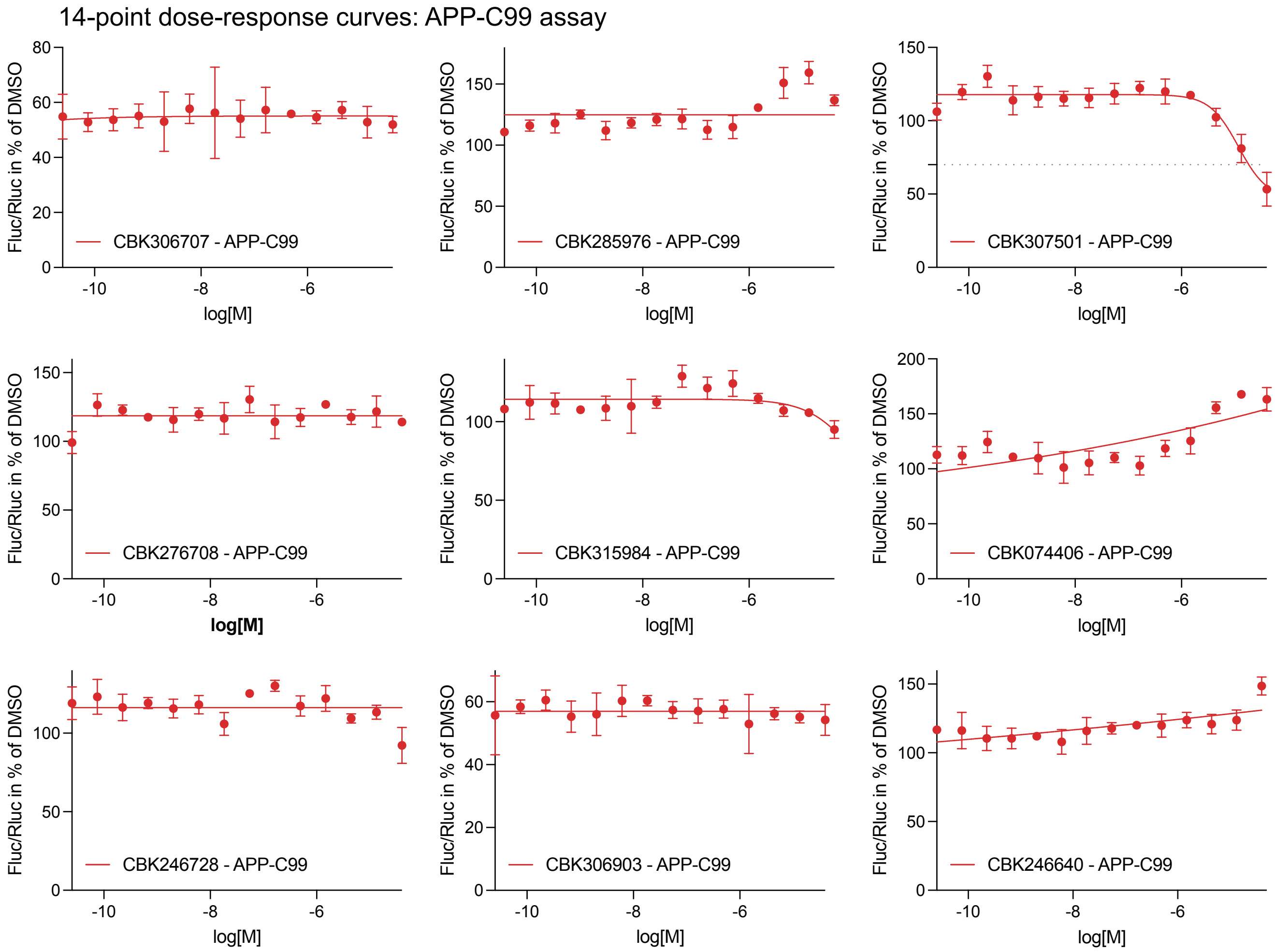

### Supplemental Figure 4C continued

14-point dose-response curves: APP-C99 assay

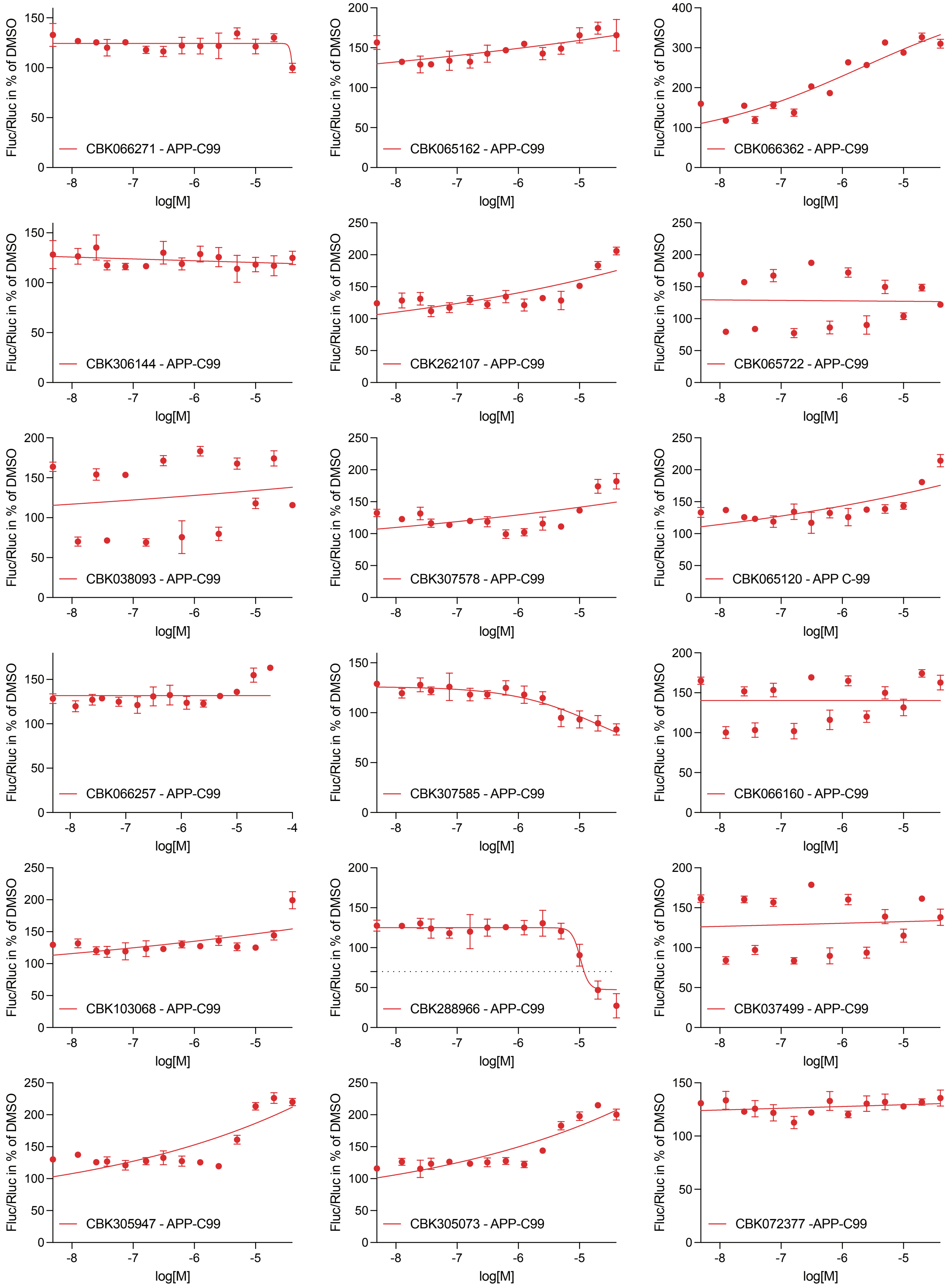

### Supplemental Figure 4C continued

14-point dose-response curves: APP-C99 assay

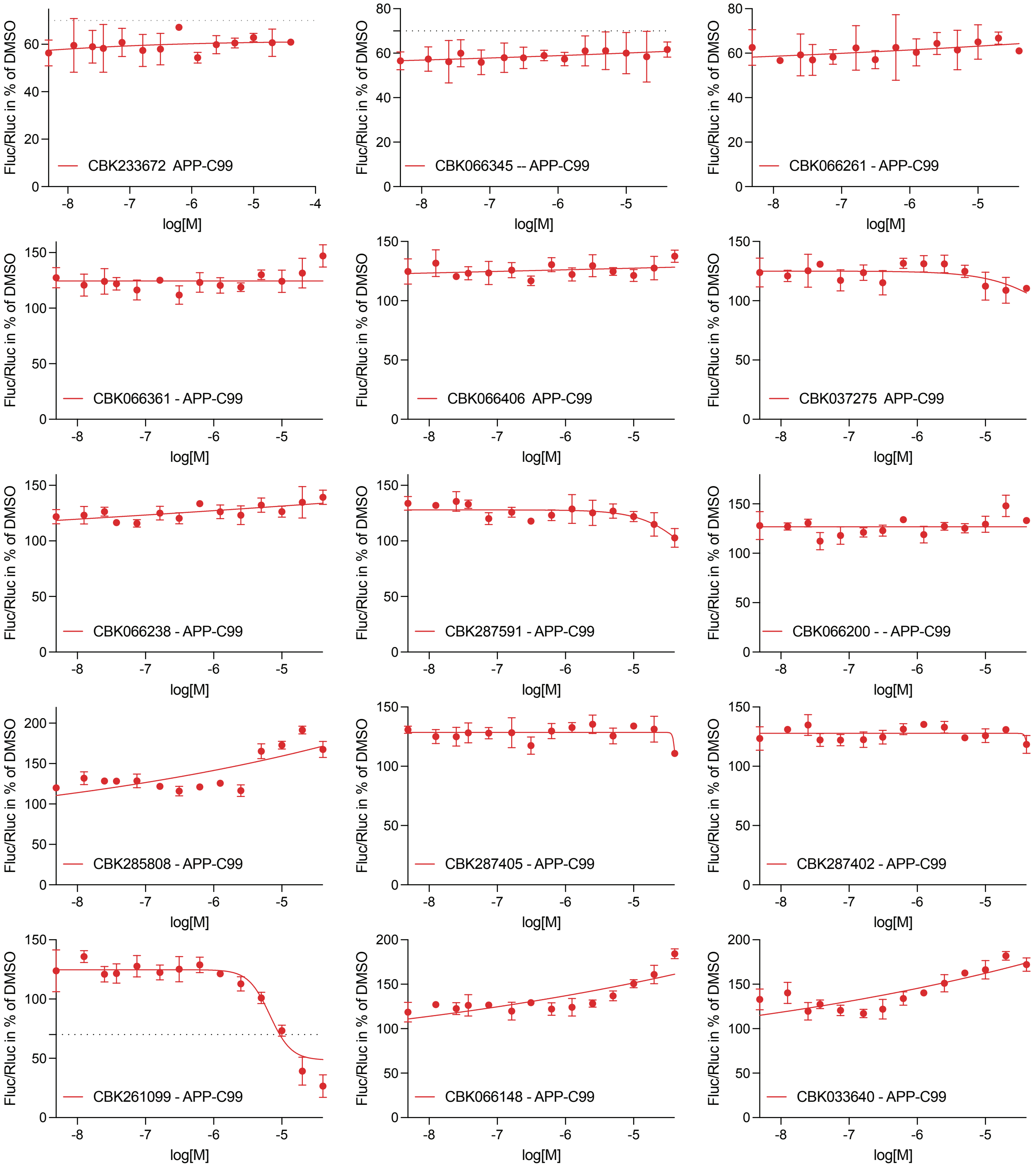

### Supplemental Figure 4C continued

#### 14-point dose-response curves: APP-C99 assay

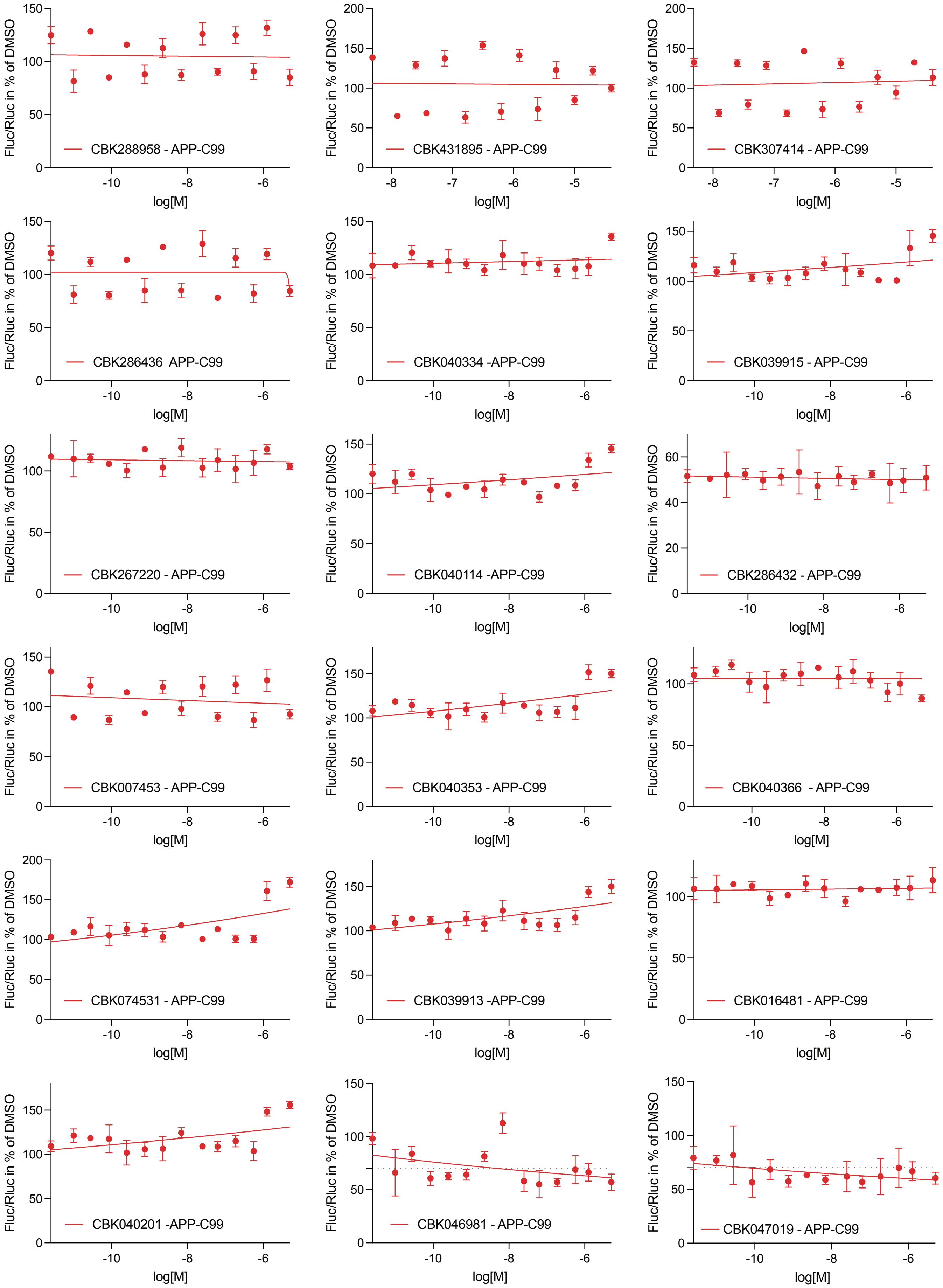

### Supplemental Figure 4C continued

14-point dose-response curves: APP-C99 assay

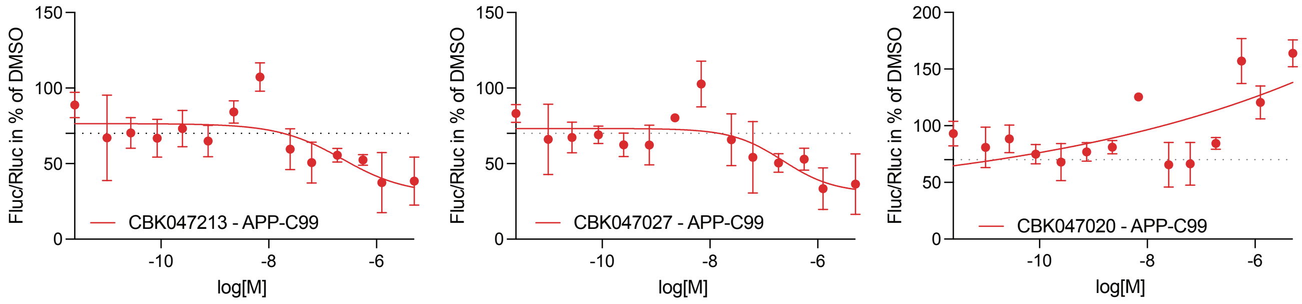

Supplemental Figure 4 continued

D

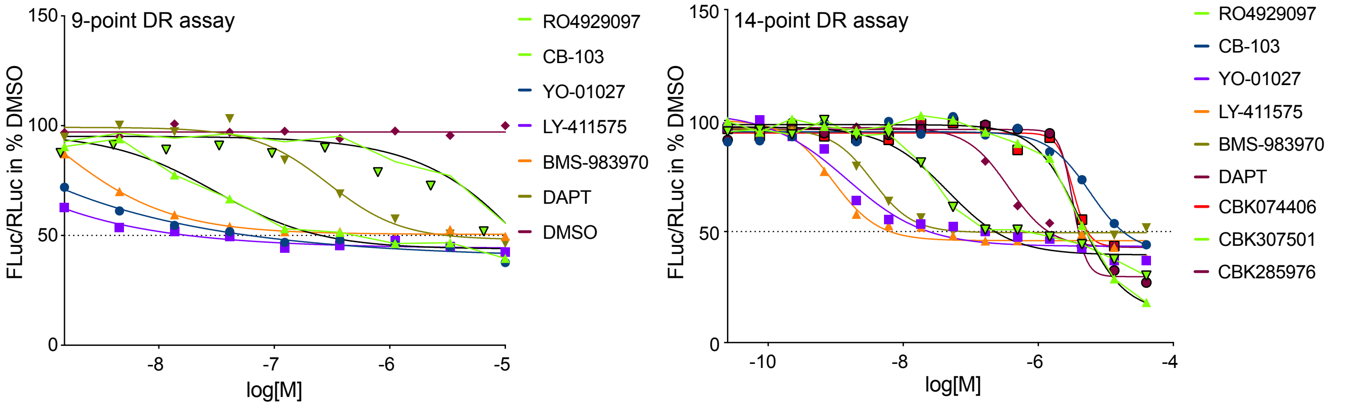


E

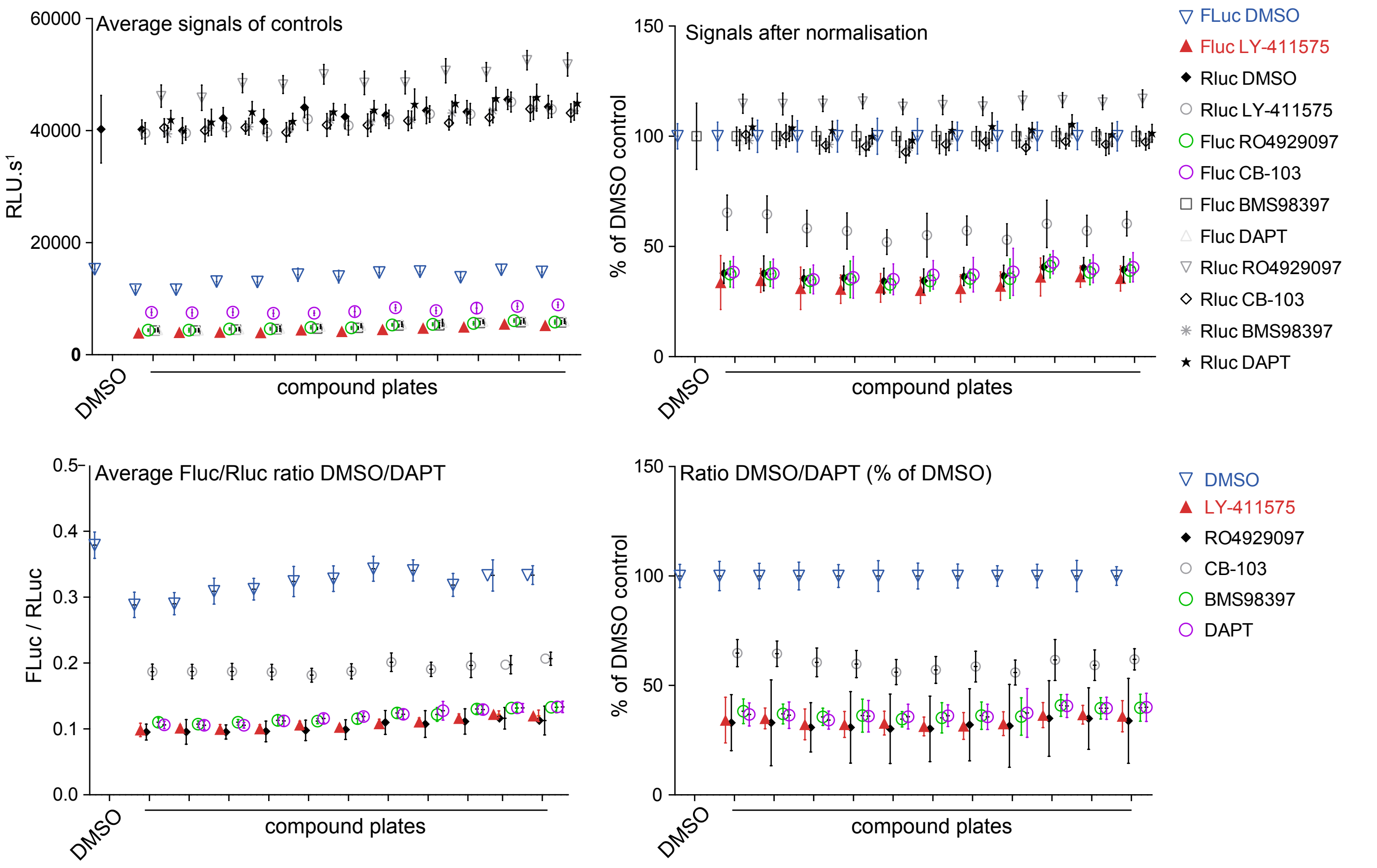


F

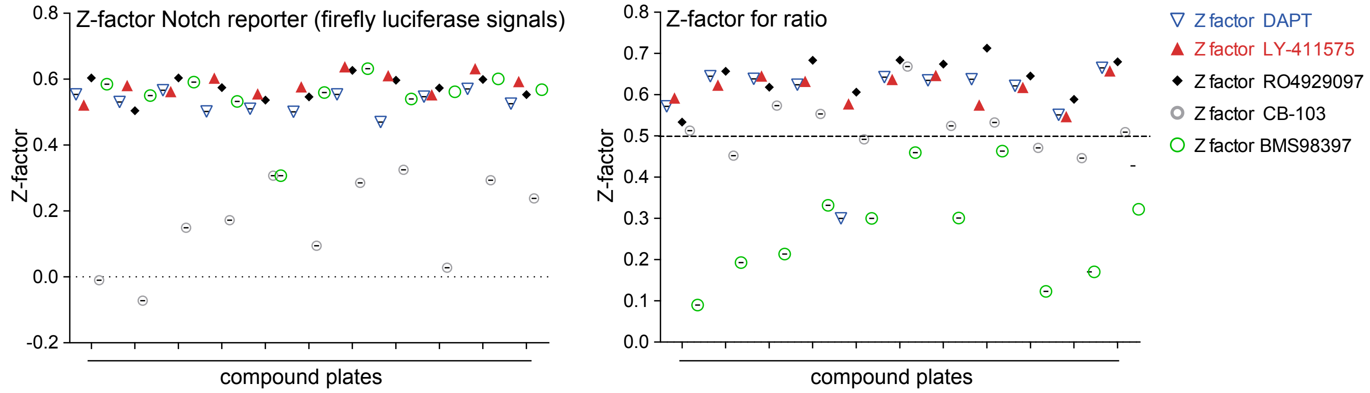

### Supplemental Figure 4 continued

G

14-point dose-response curves: Notch reporter assay

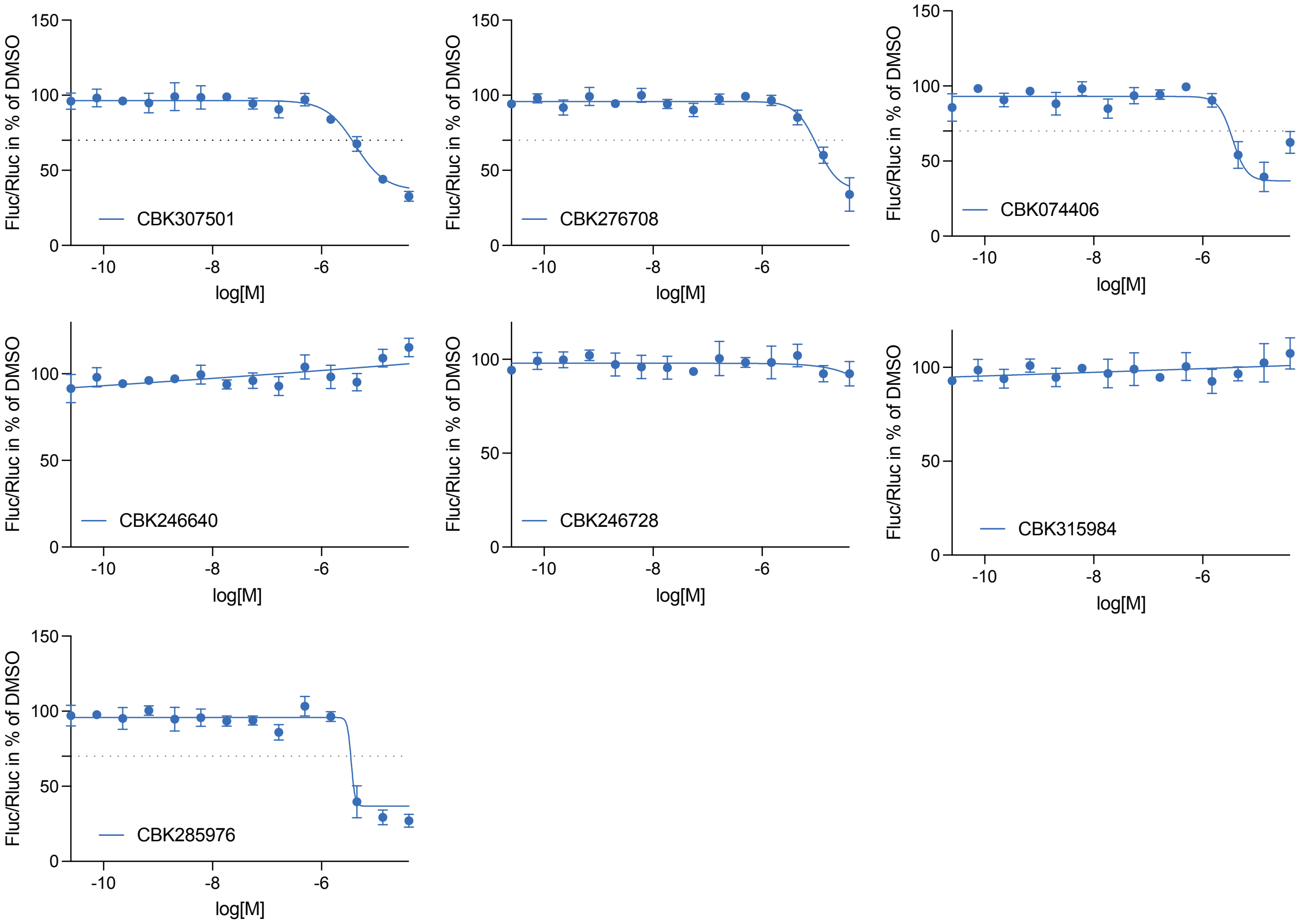

### Supplemental Figure 4G continued

14-point dose-response curves: Notch reporter assay

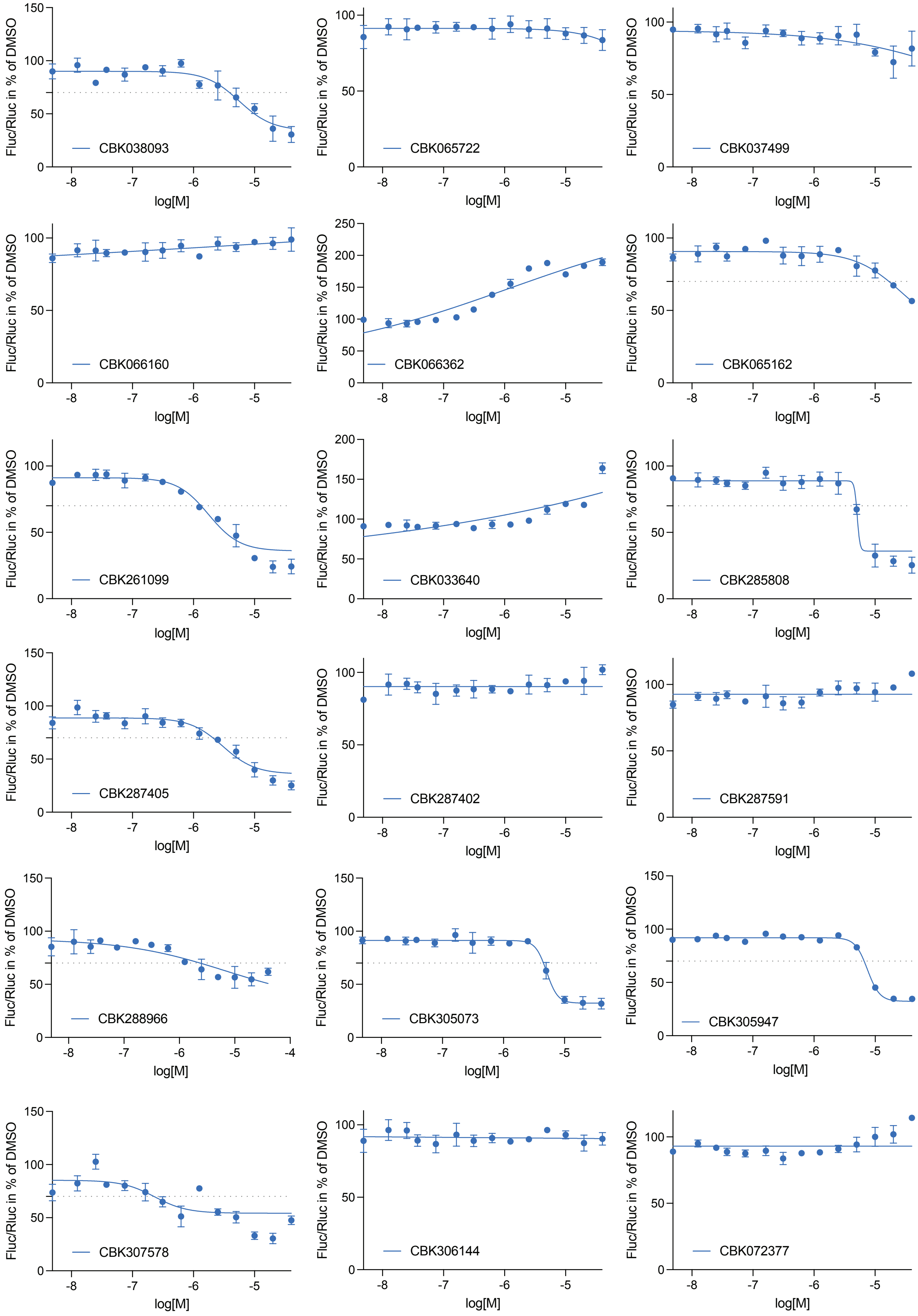

### Supplemental Figure 4G continued

14-point dose-response curves: Notch reporter assay

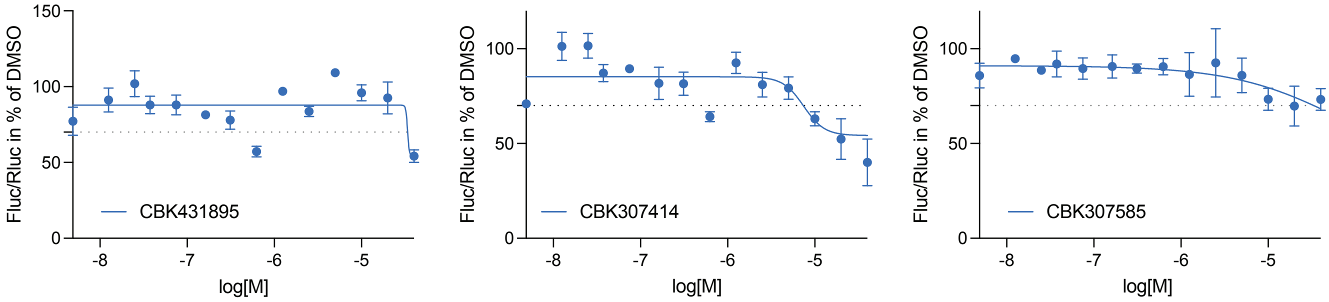

### Supplemental Figure 4G continued

14-point dose-response curves: Notch reporter assay

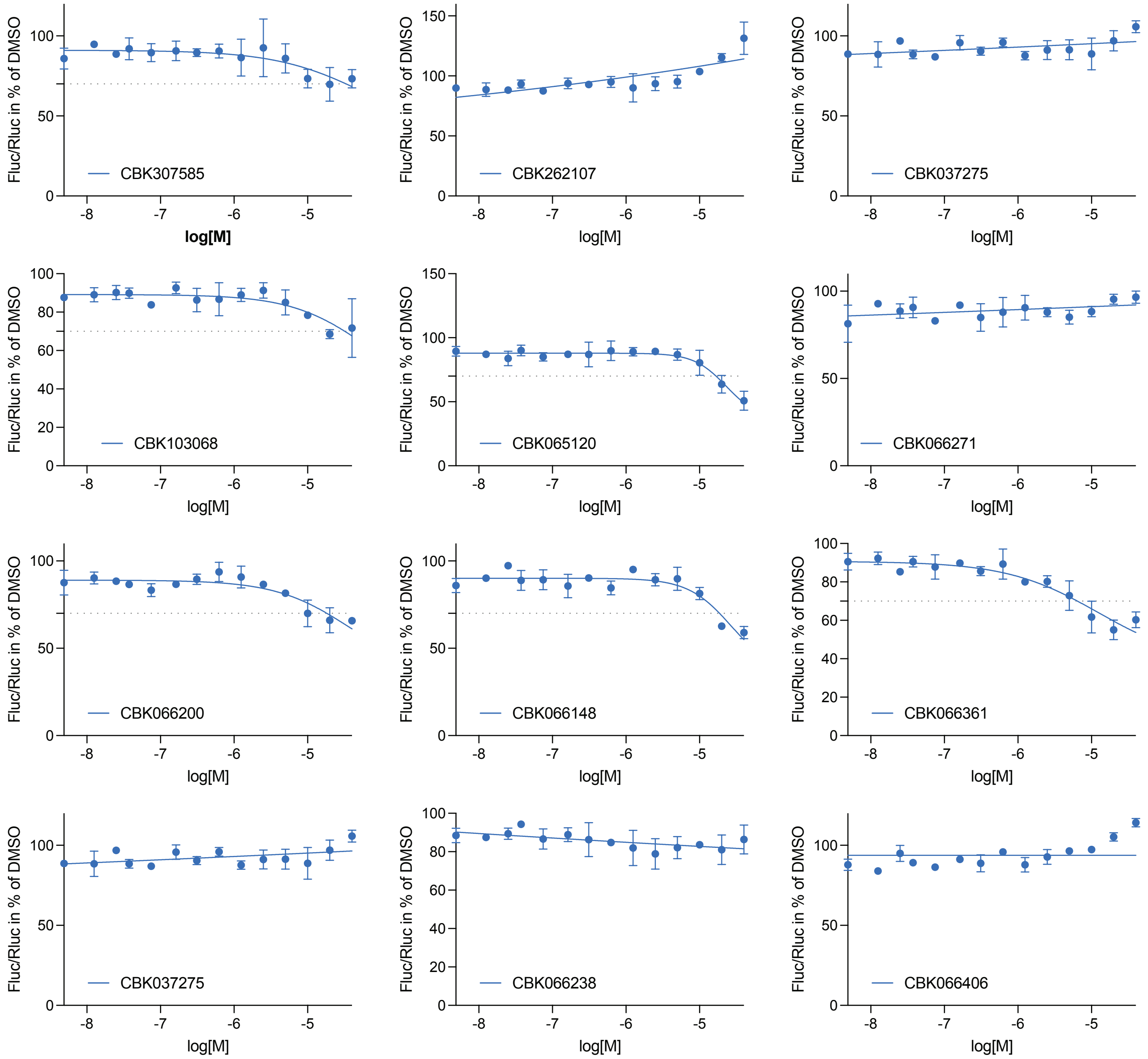

### Supplemental Figure 4G continued

14-point dose-response curves: Notch reporter assay

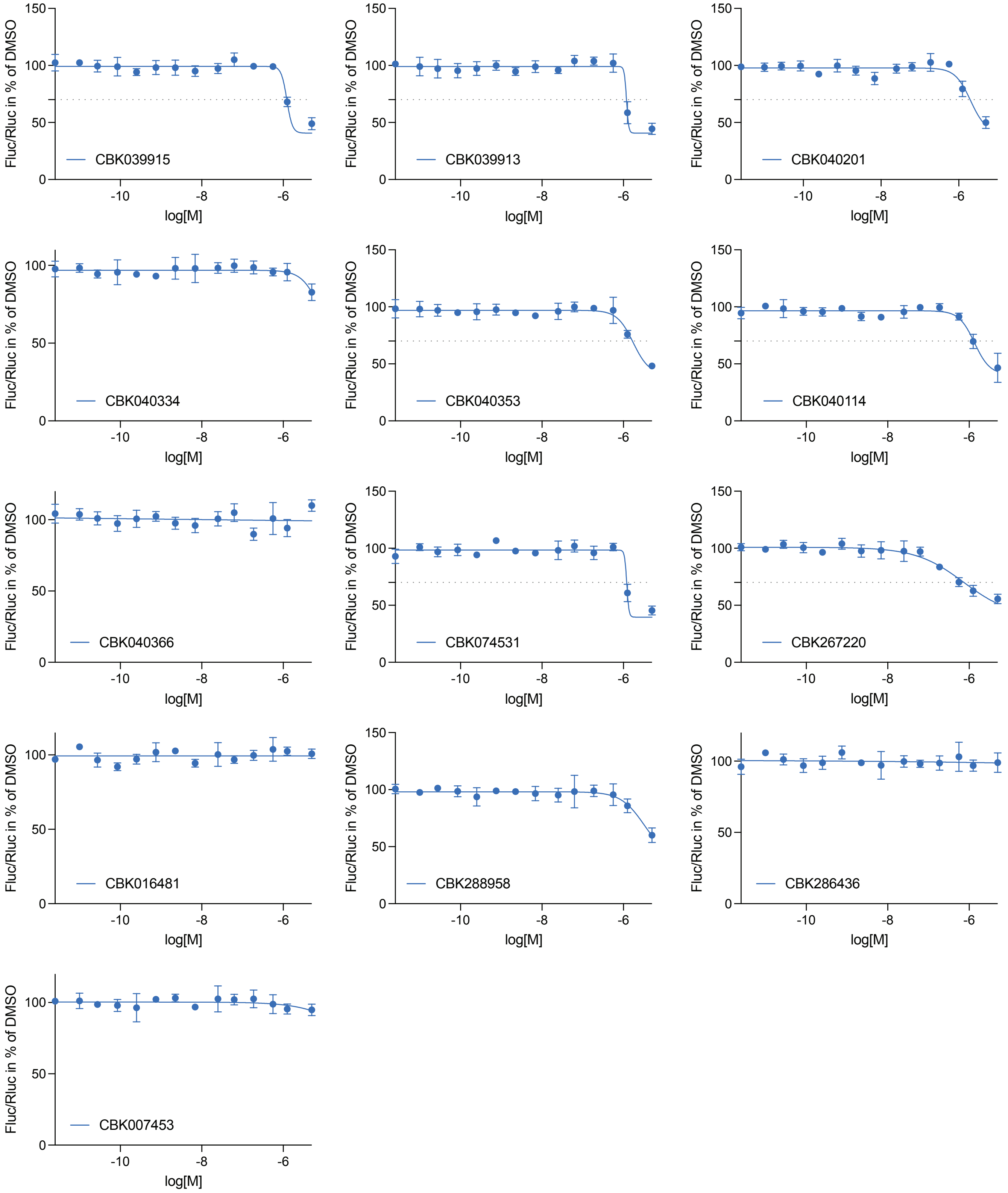

### Supplemental Figure 5

## A

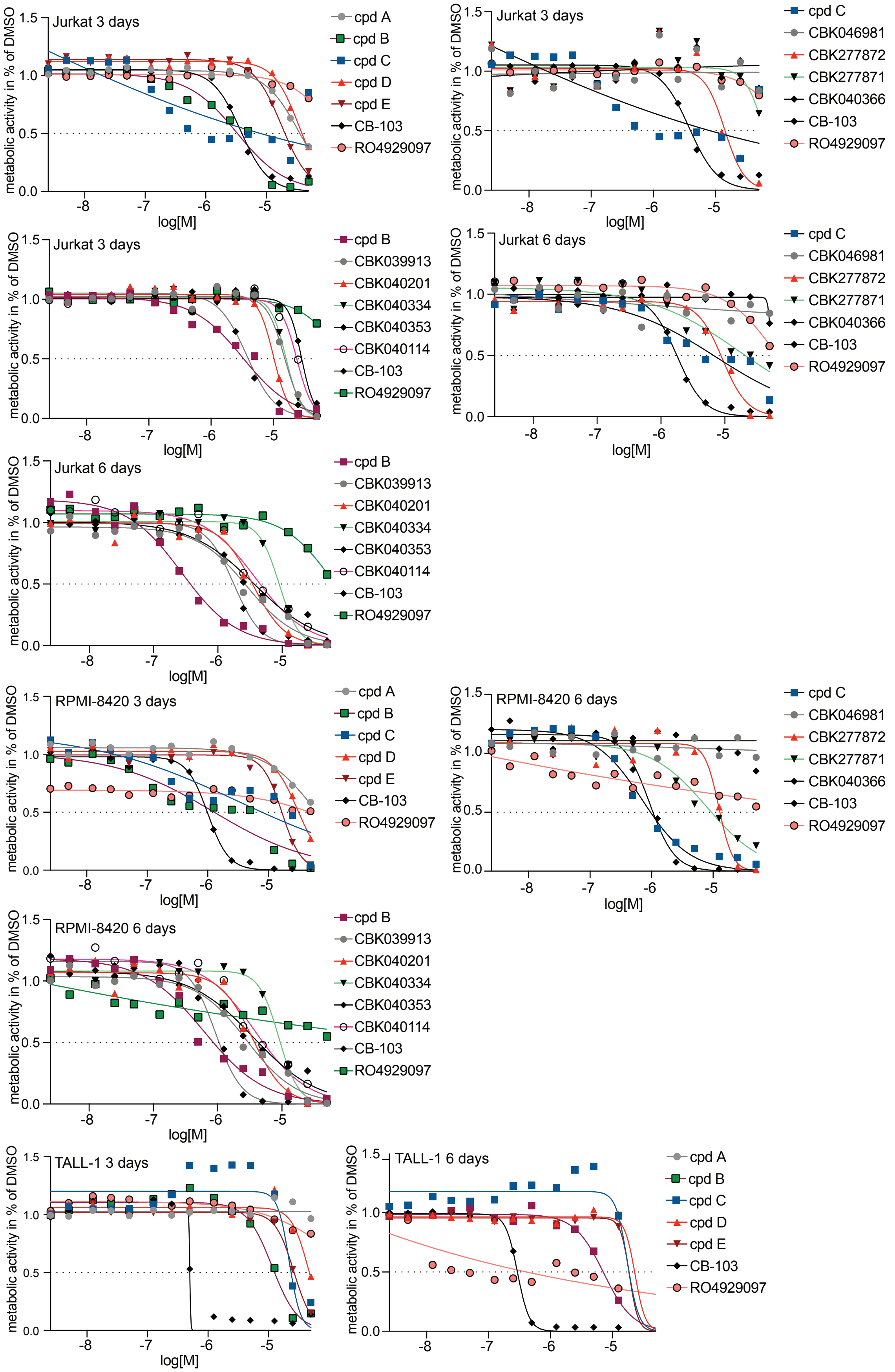

### Supplemental Figure 5 continued

B

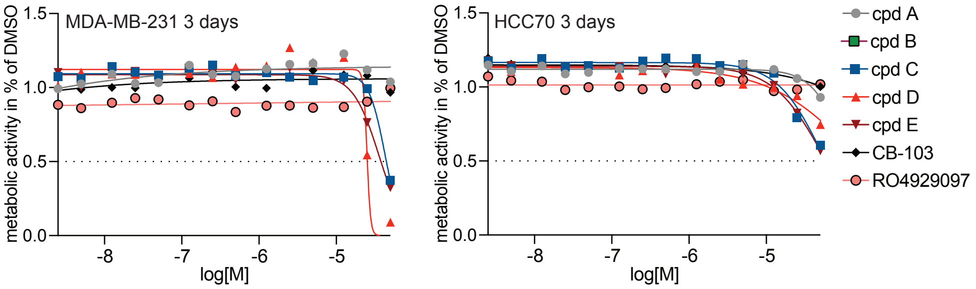


C

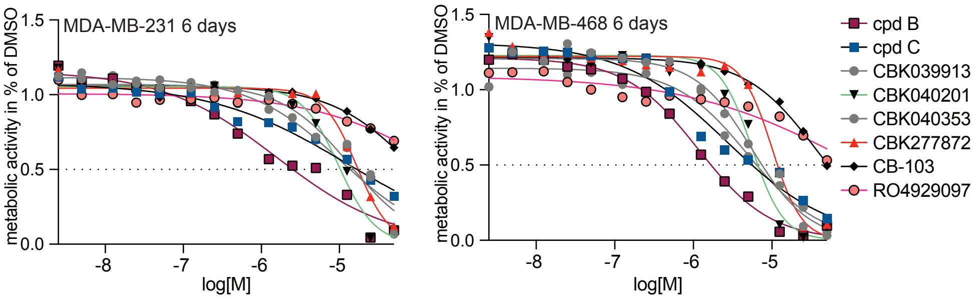

### Supplemental Figure 6

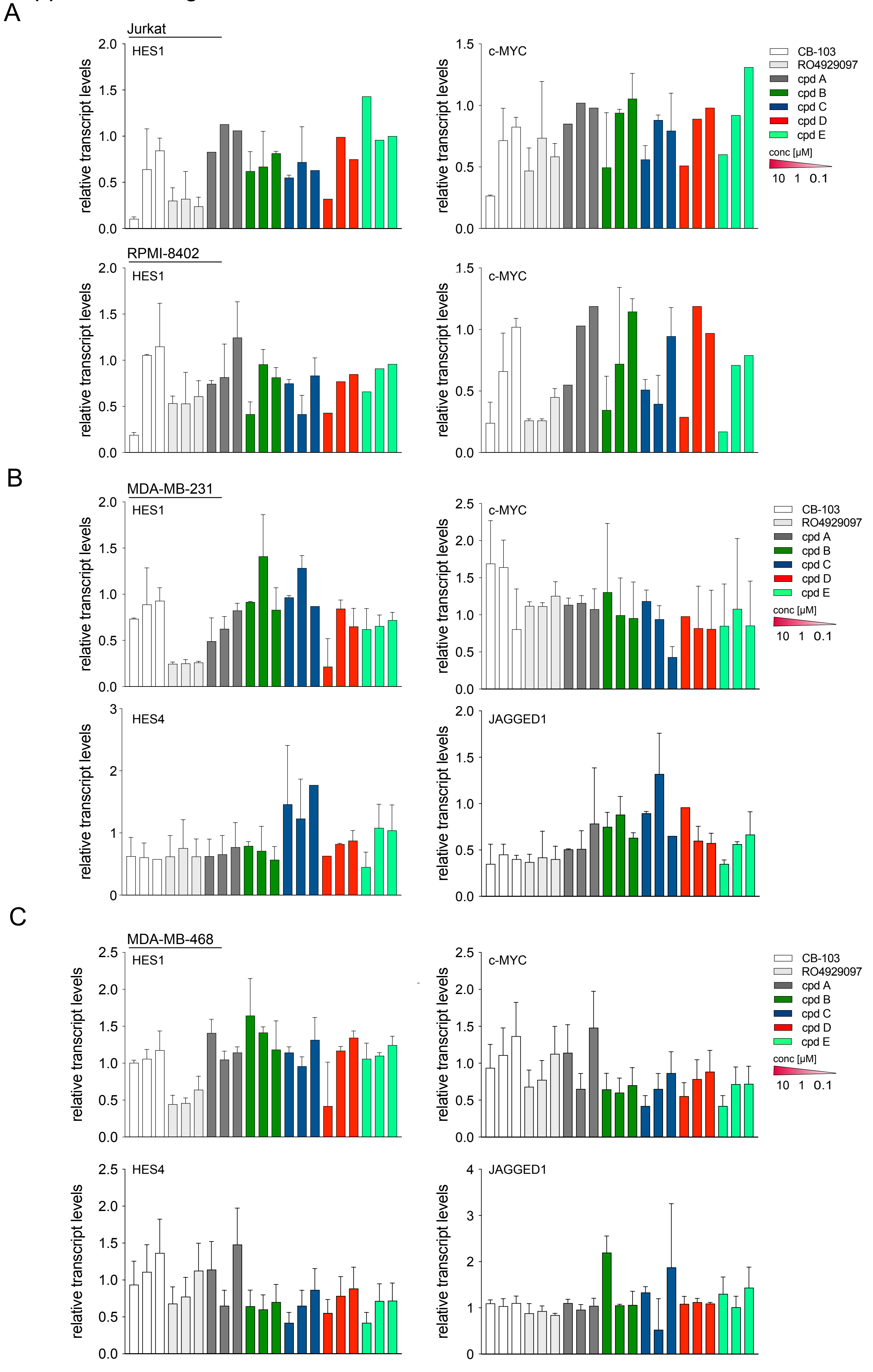


Plots with error bars represent mean ± SD of two biological replicates. Bars without error bars indicate that only technical replicates (n=1) were available

### Supplemental Figure 7

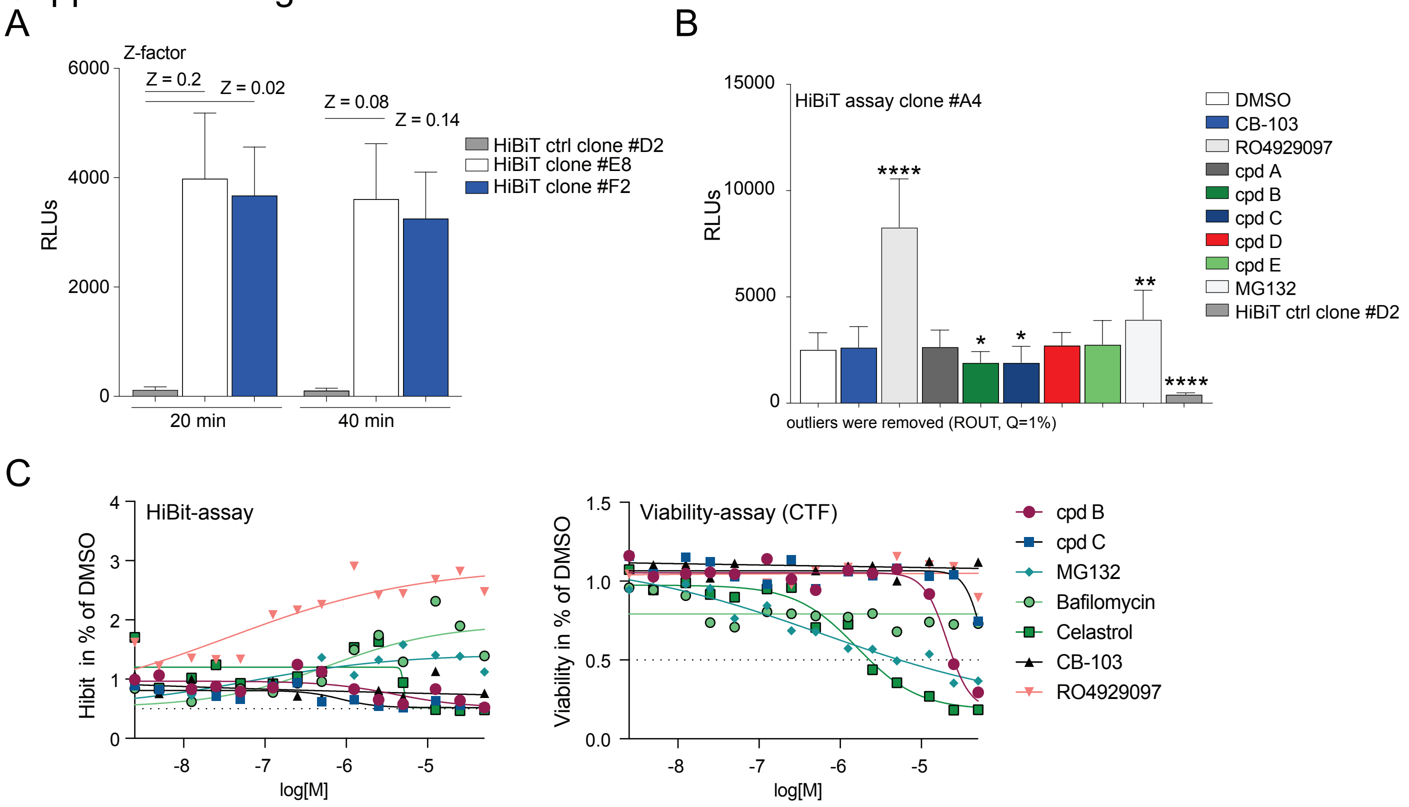
